# Rational Redesign of an Fc-Binding Peptide for Multivalent Antibody Assembly

**DOI:** 10.64898/2026.07.20.737354

**Authors:** Subhrodeep Saha, Su-Jin Lee, Hyun-Jo Shim, Song Yee Jang, Kwang-Hyun Park, Ho-Chul Shin, Eui-Jeon Woo, Jayanta Chatterjee

**Author notes:** Corresponding authors: Ho-Chul Shin, <u>Eui-jeon Woo (</u>) & Jayanta Chatterjee. These authors contributed equally to this work (Subhrodeep Saha, Su-Jin Lee).

## Abstract

Multivalent antibody assemblies offer opportunities to enhance avidity, organize immune complexes, and modulate higher-order protein interactions, but constructing such architectures from existing immunoglobulin G (IgG) molecules without redesigning the antibody scaffold remains challenging. Here, we report the rational redesign of a Protein A-derived Fc-binding peptide into ADP1, a stable dimeric Fc-binding peptide that directs Fc-mediated antibody assembly. ADP1 was designed from the parent Fc-binding peptide Z34C by preserving the Fc-recognition surface while redesigning the opposite helical surface to promote peptide–peptide association. Biophysical characterization showed that ADP1 retained nanomolar Fc-binding affinity while exhibiting markedly enhanced chemical and proteolytic stability compared with the parent peptide. Structural analyses of ADP1–Fc complexes revealed that ADP1 bridges neighboring Fc regions through a combined ADP1–Fc and ADP1–ADP1 interface, generating spiral higher-order Fc assemblies. This assembly principle was further extended to full-length IgG, where ADP1 promoted higher-order antibody association in a concentration-sensitive manner. In addition, covalent ADP1 functionalization enabled Fc-directed modification of full-length IgG while retaining Fab-mediated antigen recognition, demonstrating the utility of ADP1 as an antibody assembly and functionalization module. Finally, competitive addition of the parent Z34C peptide modulated ADP1-driven antibody assembly, suggesting a potential route for tuning Fc-mediated assembly propagation. Together, this work establishes a redesigned Fc-binding peptide platform for directing multivalent antibody assembly and functionalization without genetic reengineering of the IgG scaffold.

**Graphical abstract:** 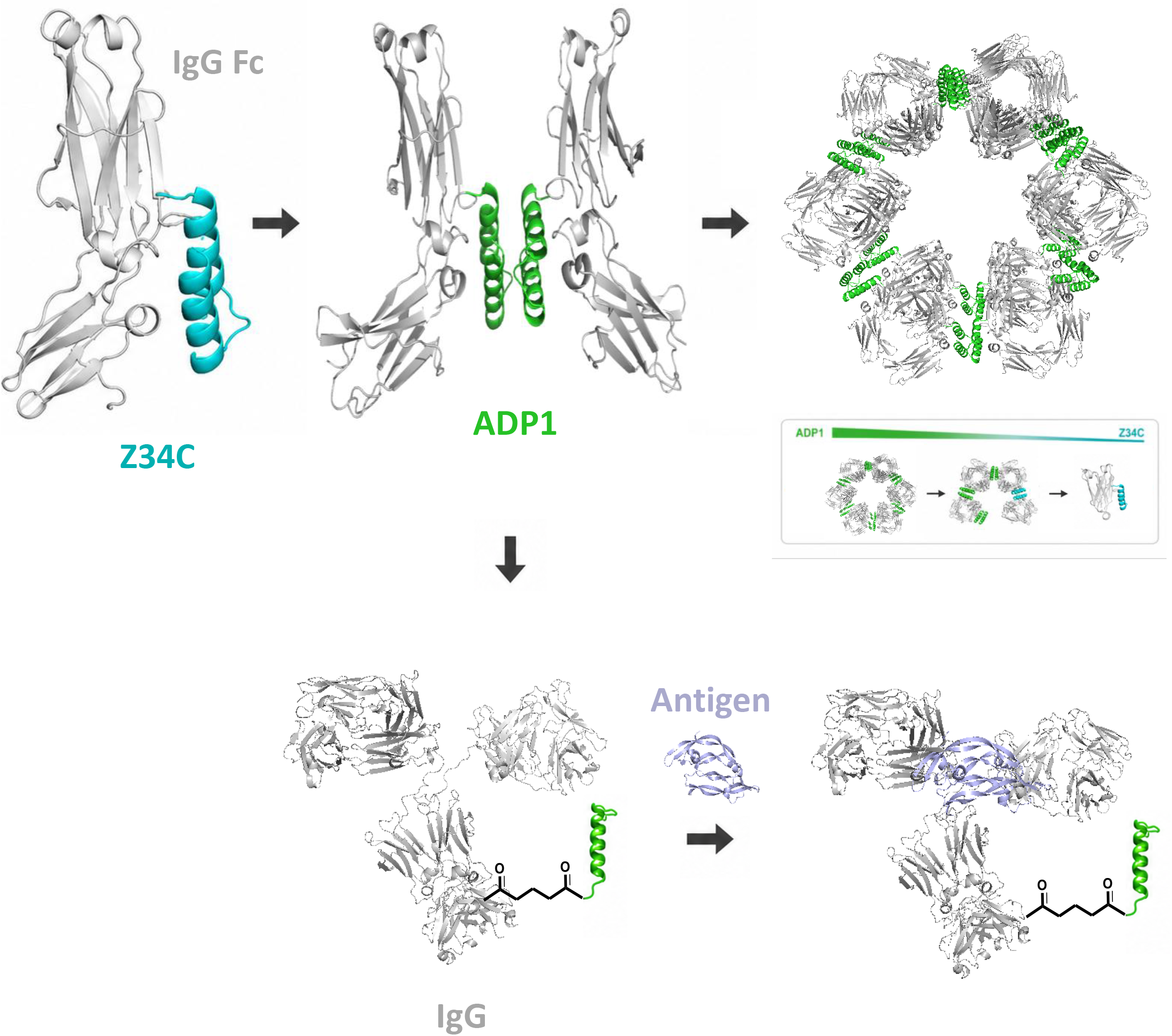

## 1. Introduction

Antibodies are among the most widely used molecular platforms in biotechnology and medicine because of their high target specificity, modular domain architecture, and established developability as biologic drugs. Although conventional immunoglobulin G (IgG) molecules function as bivalent antigen binders, naturally multivalent antibody isotypes, such as pentameric IgM, illustrate how antibody valency can influence antigen engagement^1^. This principle has motivated efforts to engineer antibody architectures with valencies and spatial organizations beyond those of native IgG^2^. However, constructing such architectures from existing IgG molecules without redesigning the antibody scaffold remains challenging.

Current strategies for generating multivalent or multifunctional antibody architectures often rely on genetic fusion, antibody reformatting, chemical conjugation, or external scaffold construction. These approaches have produced powerful formats, including bispecific antibodies, multivalent antibody constructs, and scaffolded antibody assemblies^2^. However, they can require antibody-specific engineering and may affect expression, stability, manufacturability, or the native geometry of IgG. A modular strategy that organizes existing, unmodified IgG molecules after antibody production would therefore provide a useful alternative for expanding antibody architecture while preserving the original antibody scaffold.

The fragment crystallizable (Fc) region of IgG provides an attractive handle for such modular organization. Unlike the antigen-binding Fab domains, the Fc region is relatively conserved and can be recognized by Fc-binding proteins and peptides through defined molecular interfaces. Protein A-derived Fc-binding peptides, such as Z34C, provide compact binding modules that can engage IgG Fc while leaving the Fab regions available for antigen recognition^3,4^. However, naturally derived Fc-binding peptides are primarily recognition elements; they do not by themselves create connectivity between neighboring IgG molecules. To use Fc recognition as a basis for multivalent antibody assembly, an Fc-binding peptide must be redesigned to retain Fc binding while introducing an additional interaction surface capable of linking adjacent Fc regions.

Computational protein design offers a route to repurpose such binding peptides into higher-order assembly modules. By selectively preserving residues required for Fc recognition while redesigning solvent-exposed or non-Fc-binding surfaces, a monomeric Fc-binding scaffold can potentially be converted into a dimeric or multivalent module. Related dimeric proteomimetic strategies have shown that engineered multivalency can promote target-protein dimerization through bivalent engagement^5^. In contrast, applying this concept to the conserved IgG Fc region could provide a more general strategy for assembling existing antibodies, rather than dimerizing a single disease-related target. Such a strategy could enable Fc-mediated antibody assembly without modifying the IgG scaffold itself. In addition, because the assembly module binds through Fc rather than Fab, this approach may be compatible with diverse antibodies and may preserve antigen-binding function. A further extension of this concept is Fc-directed functionalization, in which the designed Fc-binding peptide serves not only as an assembly module but also as a handle for covalent antibody modification^6^.

Here, we sought to develop an Fc-binding peptide module that could use the conserved IgG Fc region as a handle for assembling existing antibodies without modifying the antibody scaffold. To this end, we redesigned the Protein A-derived Fc-binding peptide Z34C to generate ADP1, a stable dimeric Fc-binding peptide intended to preserve Fc recognition while introducing peptide-mediated connectivity between neighboring Fc regions. We used this system to test whether Fc binding could be converted into a structural basis for higher-order Fc organization and whether the same principle could be extended to full-length IgG. We further explored whether the designed Fc-binding peptide could serve as a module for Fc-directed covalent antibody functionalization while maintaining Fab-mediated antigen recognition. Finally, we examined whether the parent Z34C peptide could modulate ADP1-driven assembly through competitive Fc binding. This study provides a modular strategy for constructing multivalent antibody assemblies from existing IgG molecules without redesigning the antibody scaffold.

## 2. Results and discussion

### 2.1. Design and validation of ADP1 as a stable dimeric IgG Fc-binding peptide

To engineer an antibody-oligomerizing miniprotein, we leveraged the structural scaffold of Z34C, a disulfide-stabilized helical hairpin peptide that binds the Fc region of IgG. Owing to its helical hairpin structure, Z34C presents two large surfaces of approximately 1,700 Å^2^ each; it utilizes one face for Fc recognition while leaving the opposite “exoface” entirely solvent-exposed^3^. Because this inert exoface does not contribute directly to Fc binding or structural stability, we targeted it for rational redesign. By substituting specific polar residues on the exoface with hydrophobic residues, we designed ADP1, a self-assembling homodimeric peptide that retains Fc-binding affinity comparable to that of Z34C while establishing bivalent connectivity between separate IgG molecules.

The design of ADP1 prioritized preserving the critical Fc-binding epitope while remodeling the opposing, non-binding helical surface to drive self-association. Based on the Z34C–Fc complex structure, two Z34C peptide chains were positioned in a geometry expected to be compatible with both Fc binding and peptide dimerization. Fc-interacting residues were fixed, whereas residues on the opposite helical surface were redesigned using ProteinMPNN to form a peptide–peptide interface. The resulting sequence candidates were evaluated using AlphaFold-Multimer to identify designs predicted to maintain the Fc-binding geometry while forming a stable dimer interface. The fixed Fc-interacting residues and tied interface residues used during the design process are summarized in **Supplementary Table 1**.

To further improve the symmetry and stability of the designed dimer interface, we performed an additional round of symmetric ProteinMPNN design in which corresponding interface residues were tied between the two peptide chains. Candidate sequences were prioritized based on AlphaFold-Multimer predictions, predicted solubility, and interface geometry. Additional point-mutation assessment was used to optimize the terminal cysteine position, resulting in the selection of ADP1, a 34-residue designed Fc-binding peptide (**Figure 1**).

**Figure 1.**
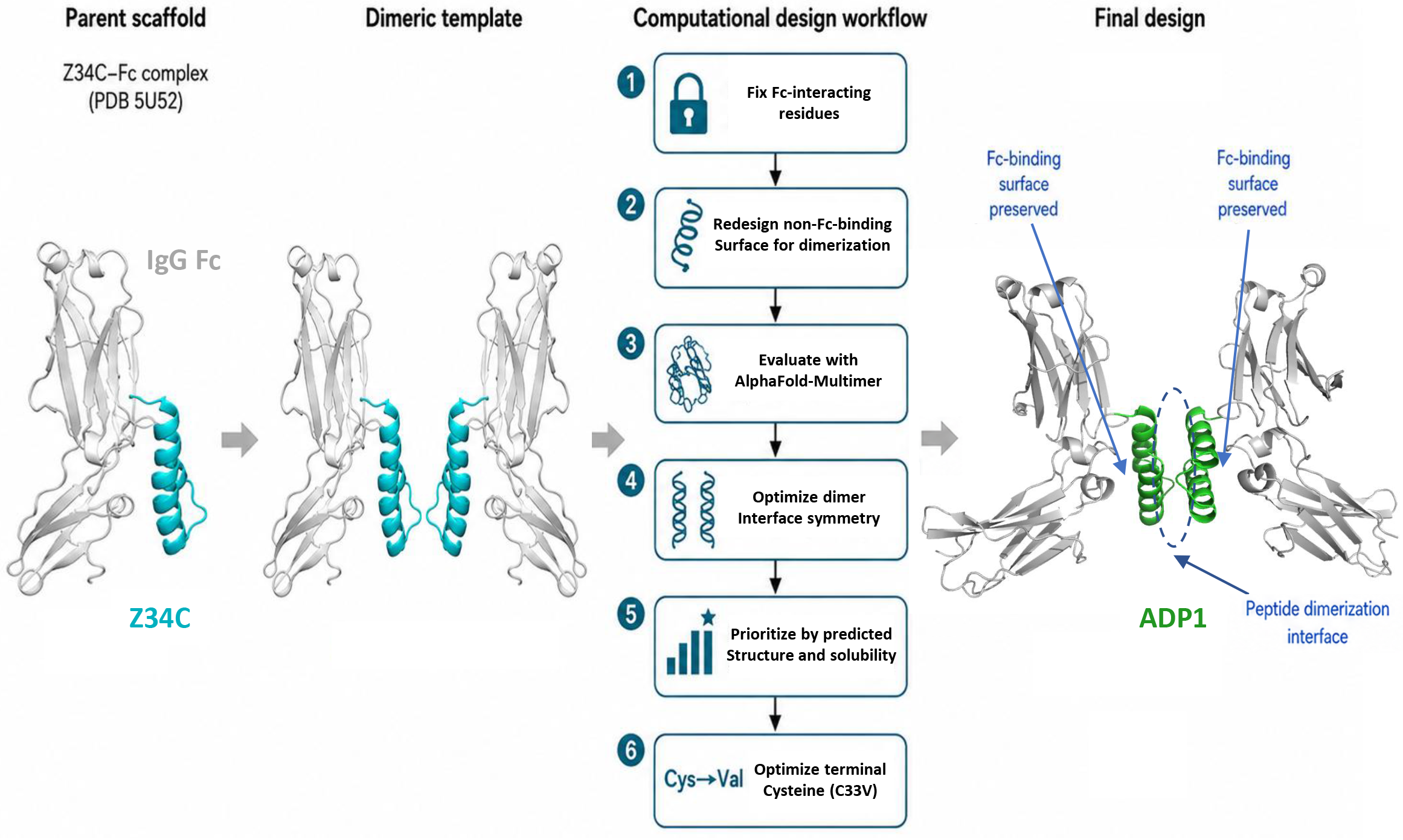
Design of ADP1 as a dimeric IgG Fc-binding peptide. Structure-guided computational design workflow of ADP1 from the Protein A-derived Fc-binding peptide Z34C.

Sedimentation velocity analytical ultracentrifugation confirmed that the parent Z34C peptide existed predominantly as a monomer, whereas ADP1 assembled into a well-defined homodimer (**Figure 2A**). Despite the extensive sequence modifications introduced to drive self-association, circular dichroism (CD) spectroscopy revealed that ADP1 retained a highly helical secondary structure comparable to that of Z34C (**Figure 2B, C**). The thermal stability, helicity, and Fc-binding affinities of Z34C, ADP1, and the ADP1-derived variants are summarized in **Supplementary Table 2.** Notably, engineered homodimerization yielded a substantial increase in thermal stability, with ADP1 exhibiting a melting temperature above 95°C, compared with 59.4°C for Z34C **(Supplementary Figure 1A)**. Homodimerization also completely restored structural reversibility. Whereas the parent Z34C peptide failed to refold after thermal denaturation, ADP1 exhibited near-complete refolding upon cooling after thermal denaturation (**Figure 2B, C)**. Dimerization also significantly accelerated the oxidative folding kinetics of the peptide, with ADP1 demonstrating a more than 6-fold increase in the rate of native disulfide bond formation relative to Z34C (**Supplementary Figure 1B**). Furthermore, whereas the native Z34C exoface is predominantly polar, the hydrophobic substitutions introduced to promote homodimerization generated a tightly packed hydrophobic core analogous to the interior of a globular protein. This engineered core provided substantial protection against chemical and enzymatic degradation, yielding a more than 50-fold increase in disulfide stability against dithiothreitol (DTT)-mediated reduction (**Figure 2D**) and a more than 10-fold increase in proteolytic resistance against Proteinase K degradation (**Figure 2E**). Together, these data demonstrate that the computational redesign of the Z34C exoface not only successfully promoted homodimer formation and bivalent connectivity but also conferred exceptional structural, chemical, and proteolytic stability on the miniprotein.

**Figure 2.**
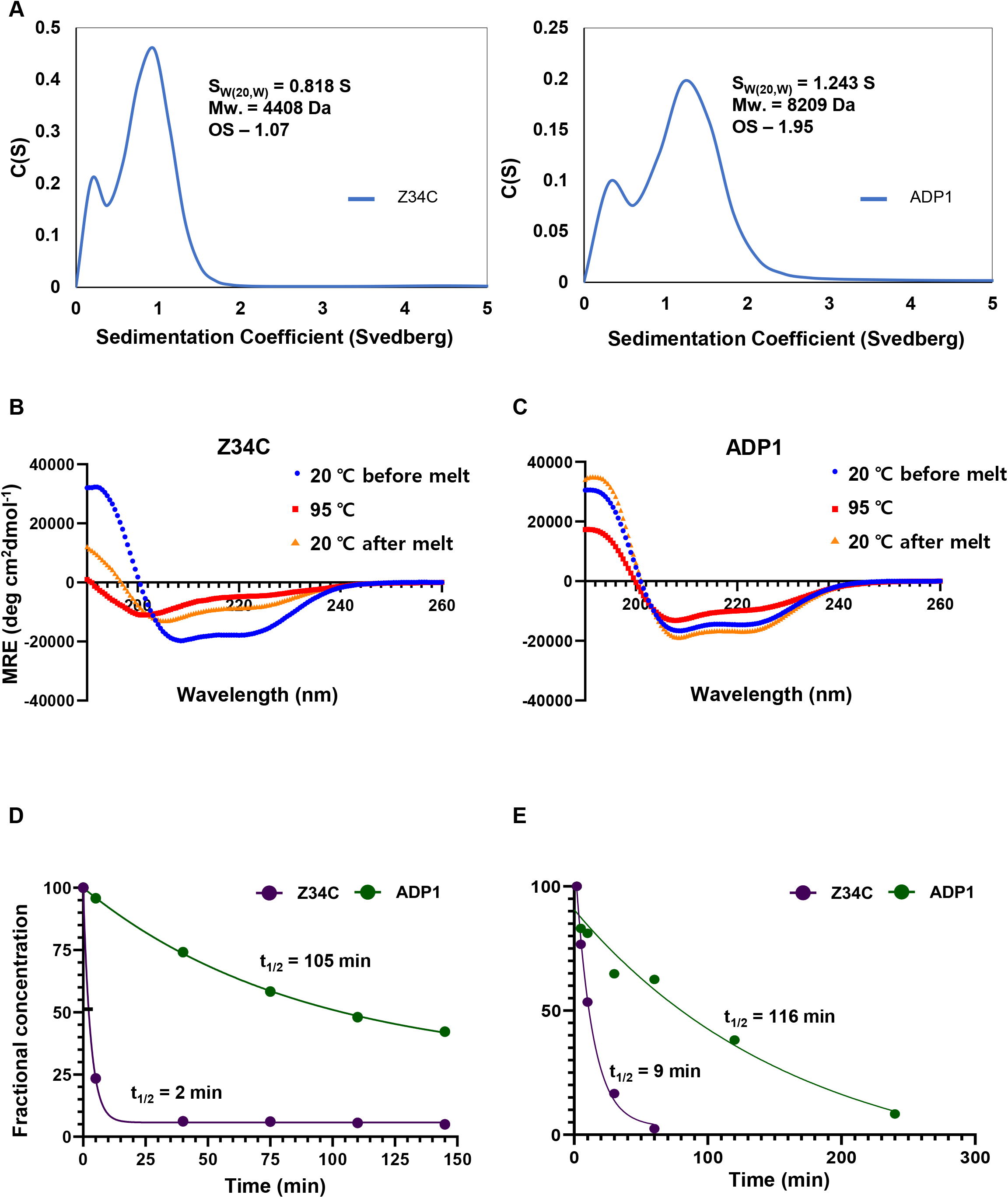
Biophysical characterization of ADP1. **(A)** AUC of Z34C and ADP1. **(B)** CD plots of Z34C and **(C)** ADP1. **(D)** Reduction kinetics of Z34C and ADP1 in 2mM DTT. **(E)** Proteolysis of Z34C and ADP1 in Proteinase K. Molecular masses shown in panel A correspond to the major sedimenting species identified in the AUC analysis.

### 2.2. Structural basis of ADP1-mediated spiral Fc assembly

Having established ADP1 as a stable, homodimeric miniprotein, we investigated its capacity to drive the higher-order assembly of individual Fc domains. We first incubated ADP1 with commercial, human serum-derived IgG Fc (purchased from Merck) and evaluated the complex by size-exclusion chromatography coupled with multi-angle light scattering (SEC-MALS). The SEC-MALS profile revealed the formation of a distinct high-molecular-weight species that eluted in the column void volume (**Supplementary Figure 2A**). For cryo-EM sample preparation, the Fc–ADP1 mixture was subsequently separated by size-exclusion chromatography, and the early-eluting assembly-associated fraction was collected and concentrated for grid preparation **(Supplementary Figure 3)**. The isolated fraction was then subjected to cryogenic electron microscopy (cryo-EM) analysis. Three-dimensional reconstruction successfully resolved a spiral helical Fc assembly at an overall resolution of 4.88 Å (**Figure 3A** and **Supplementary Table 3)**. This result indicates that ADP1 can organize Fc fragments into an ordered supramolecular architecture composed of repeating Fc-associated units, rather than forming nonspecific aggregates. In the cryo-EM structure, Fc units are arranged sequentially along the helical axis, with density corresponding to a bridging ADP1 homodimer positioned between neighboring Fc units. The assembly showed a helical pitch of approximately 299.4 Å and an outer helical diameter of 157.7 Å, with approximately 4.1 Fc-associated units per turn (**Figure 3A**). This arrangement is consistent with our design principle that ADP1 functions not merely as an Fc-recognition module, but as a structural linchpin that couples target binding to programmed macromolecular assembly. Crucially, closer inspection of the reconstruction revealed significant structural flexibility near the N-terminal hinge region, where the individual Fc protomers within the homodimer appeared to separate near the hinge. We hypothesized that this separation potentially stemmed from a lack of intact inter-chain disulfide bonds within the commercial, serum-derived pool, likely disrupted during industrial processing or purification.

**Figure 3.**
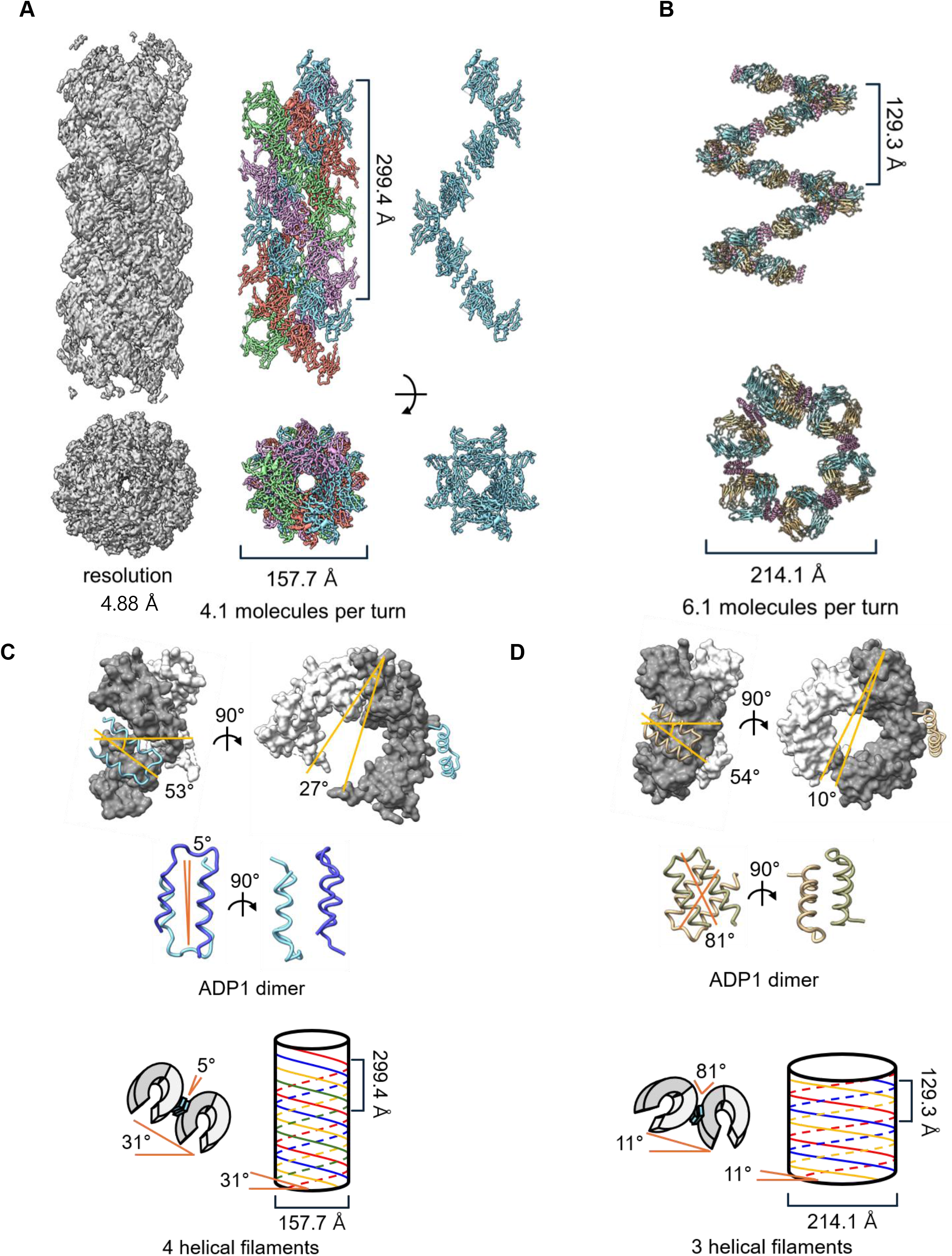
Structural basis of ADP1-mediated spiral Fc assembly. **(A)** Cryo-EM structure of the ADP1-induced Fc assembly. **(B)** X-ray crystal structure of the ADP1-bound bevacizumab Fc complex. **(C)** Helical arrangement and ADP1 dimer geometry derived from the cryo-EM structure. **(D)** Helical arrangement and ADP1 dimer geometry derived from the X-ray structure.

To resolve this structural bottleneck, we investigated whether ADP1 could drive a more rigid, higher-order assembly when paired with a homogeneous, monoclonal antibody-derived IgG1 Fc domain. Standard papain cleavage protocols typically require mild reducing conditions (10-20 mM cysteine) to activate the protease; however, this environment routinely reduces the native inter-chain disulfide bonds of the antibody, yielding a heterogeneous mixture of Fc fragments with structurally compromised hinges^7,8^. To circumvent this, we optimized a papain-mediated digestion protocol using the monoclonal antibody bevacizumab under strict non-reducing conditions, precisely limiting the incubation duration to prevent over-digestion of the polypeptide chain below the core hinge region.^9,10^ By generating a homogeneous preparation of IgG1 Fc fragments designed to retain these native inter-chain disulfide bridges, we sought to mitigate hinge-region flexibility during subsequent structural characterization. To this end, we next evaluated the assembly profile of ADP1 when incubated with this in-house-generated bevacizumab Fc using SEC-MALS. In contrast to a broad high-molecular-weight assembly fraction observed with the human serum-derived pool, SEC-MALS analysis of the bevacizumab Fc–ADP1 complex resolved distinct, well-defined macromolecular species corresponding to discrete higher-order oligomeric states. Two higher-molecular-weight species were observed at approximately 139 and 216 kDa, consistent with discrete Fc–ADP1 oligomeric assemblies (**Supplementary Figure 2B**). To elucidate the high-resolution structural details governing this assembly, we determined the X-ray crystal structure of the ADP1–bevacizumab Fc complex at a resolution of 2.6 Å (**Figure 3B** and **Supplementary Table 4)**. The X-ray structure confirmed at high resolution that ADP1 binds Fc while preserving the Protein A-derived Fc-binding mode of the parent Z34C peptide^4^. Specifically, ADP1 engaged the Fc homodimer through its Fc-recognition surface, whereas the opposite redesigned exoface formed a peptide–peptide interface with an adjacent ADP1 chain. This arrangement directly validates our design strategy, wherein the solvent-exposed, polar exoface of the parent Z34C scaffold was converted into an assembly-driving hydrophobic core via targeted polar-to-hydrophobic substitutions **(Supplementary Figure 4A, B)**. In addition, overlay of the designed ADP1–Fc model with the experimentally determined ADP1–Fc X-ray structure showed close agreement in the Fc-bound conformation, with an RMSD of 0.46 Å, supporting the structural accuracy of the computational design **(Supplementary Figure 4C)**.

The cryo-EM reconstruction and crystallographic symmetry expansion of the X-ray structure both showed that ADP1 organizes Fc regions into spiral higher-order assemblies, although the two assemblies differed in geometry. The cryo-EM and crystallographic assemblies also differed in higher-order packing, comprising four and three associated helical filaments, respectively. The X-ray-derived assembly had a helical pitch of approximately 129.3 Å and a helical width of 214.1 Å, with approximately 6.1 Fc-associated units per turn (**Figure 3B**). Thus, the cryo-EM assembly displayed a relatively longer pitch and narrower width, whereas the X-ray assembly showed a shorter pitch and wider helical organization. The geometry analyses in **Figure 3C**, **D** further indicate that these differences arise from changes in the relative orientation of neighboring Fc regions and the arrangement of the ADP1 dimer. In both structures, ADP1 binds the Fc surface and connects adjacent Fc regions through the ADP1–ADP1 interface. However, in the cryo-EM structure, the ADP1 dimer is arranged with a relatively small crossing angle along the assembly axis, whereas in the X-ray structure, the ADP1 dimer adopts a larger crossing angle (**Figure 3C, D)**. This difference in the relative arrangement of the ADP1–ADP1 interface is also evident in **Supplementary Figure 5B**. Crucially, in stark contrast to the pronounced structural separation observed near the N-terminal hinge of the serum-derived Fc in the cryo-EM map, the bevacizumab Fc protomers within the crystal structure did not exhibit comparable separation. This structural observation strongly supports our hypothesis that preservation of the native inter-chain disulfide bonds effectively restrains the hinge conformation. Together, the two structures suggest that the same ADP1-mediated connectivity can propagate into distinct helical trajectories depending on Fc conformation and assembly environment. Factors such as hinge and inter-chain disulfide status, the degree of Fc opening, glycan or chain heterogeneity, sample preparation, and crystal-packing constraints may influence the relative orientation of neighboring Fc regions.

The molecular interactions stabilizing ADP1-mediated assembly further support this structural interpretation. At the ADP1–Fc interface, π–π interactions, polar contacts, salt bridges, and hydrophobic contacts were observed, supporting preservation of the Protein A-derived Fc-binding mode **(Supplementary Figure 5A)**. At the ADP1–ADP1 interface, redesigned hydrophobic residues and salt-bridge interactions contributed to peptide dimerization **(Supplementary Figure 5B)**. Thus, the ADP1 homodimer functions as a bifunctional, bivalent Fc-connecting module: each peptide chain engages Fc through its preserved Fc-recognition surface, while the redesigned exoface mediates ADP1–ADP1 association.

Together, the cryo-EM and X-ray structures provide complementary structural evidence for ADP1-mediated Fc assembly. The cryo-EM structure shows that ADP1 can organize human serum-derived IgG Fc into a spiral higher-order architecture in a solution-derived assembly fraction, whereas the X-ray structure provides the atomic basis of ADP1–Fc recognition and ADP1–ADP1 dimerization. Although the two structures differ in helical pitch, assembly width, molecules per turn, and ADP1 dimer geometry, both support the common principle that ADP1 converts Fc recognition into peptide-mediated Fc-to-Fc connectivity. These results establish ADP1 as an assembly-driving Fc connector capable of linking Fc recognition to higher-order antibody assembly.

### 2.3. ADP1-mediated higher-order assembly of full-length IgG

The Fc-level structures suggested that ADP1 could connect neighboring Fc regions through peptide-mediated dimerization. However, transitioning from these truncated fragments to full-length IgG introduces a significantly more complex architecture, where the flexible Fab arms could potentially impose severe steric constraints on Fc-mediated organization. We therefore sought to determine whether this design principle remains viable in the context of an intact antibody.

Analytical ultracentrifugation analysis showed that bevacizumab alone was predominantly monomeric, with an apparent molecular weight consistent with full-length IgG. In contrast, bevacizumab incubated with ADP1 showed a broader distribution containing species larger than monomeric IgG, indicating that ADP1 can promote antibody association in the context of full-length IgG, yielding an oligomeric distribution with a population centered at approximately 701 kDa (**Figure 4A**). The higher apparent molar mass observed under the high-concentration SEC-MALS conditions is consistent with the possibility that higher protein concentrations favor larger assemblies, although direct comparison is limited by differences between the two analytical methods and measurement conditions. Crucially, because AUC evaluates macromolecular sedimentation directly in free solution without the surface interactions of a chromatographic matrix, the detection of these larger complexes demonstrates that ADP1-mediated assembly occurs in free solution and is not solely attributable to column-induced effects. These results suggest that the Fc-to-Fc connectivity observed in Fc-level structural analyses can be transferred to intact antibodies without genetic modification of the antibody scaffold.

**Figure 4.**
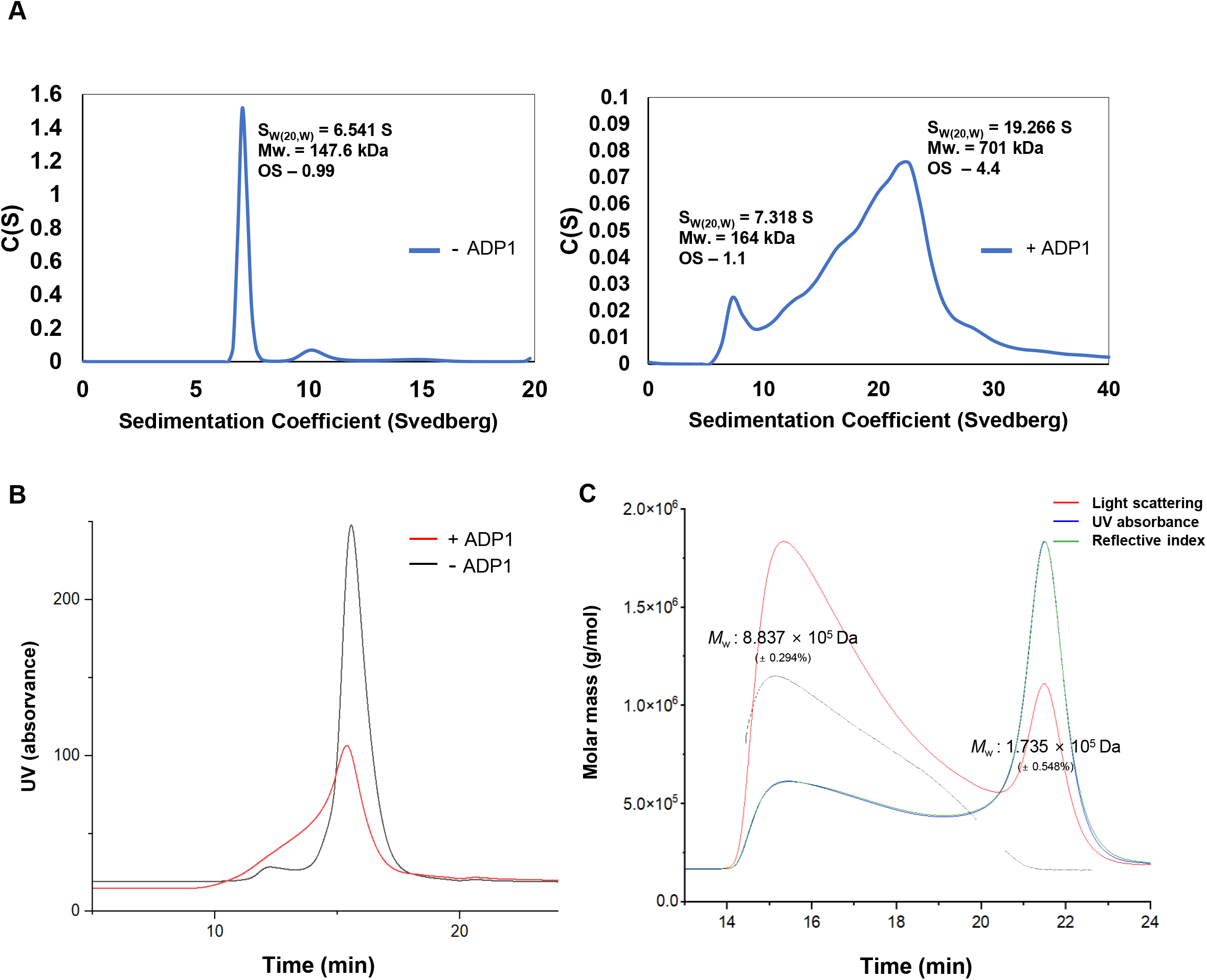
ADP1-mediated higher-order assembly of full-length IgG. **(A)** Analytical ultracentrifugation analysis of bevacizumab in the absence or presence of ADP1. **(B)** SEC analysis of bevacizumab in the absence or presence of ADP1 under high-concentration conditions. **(C)** SEC–MALS analysis of bevacizumab incubated with ADP1 under high-concentration conditions. Molecular masses shown in panel A correspond to the major sedimenting species identified in the AUC analysis.

We further examined ADP1-mediated full-length IgG assembly under high-concentration conditions that exceed the operational limits of optical detection in AUC^11^, where Fc-mediated propagation would be expected to become more apparent. Size-exclusion chromatography analysis showed that bevacizumab incubated with ADP1 produced an additional early-eluting species compared with bevacizumab alone, consistent with the formation of higher-order antibody assemblies (**Figure 4B**). SEC-MALS analysis resolved a higher-order species with a measured (**Figure 4C**).

Together, the AUC, SEC, and SEC–MALS data provide complementary evidence that ADP1 can promote higher-order assembly of full-length IgG through Fc-mediated connectivity. Importantly, this assembly was achieved using intact, unmodified antibodies without any genetic modification of the IgG scaffold, indicating that the Fc-directed design principle observed in Fc-fragment structures can be translated to full-length antibody systems. These results establish ADP1 as a minimal post-production peptide module capable of organizing existing therapeutic IgG antibodies and provide the basis for extending the platform toward Fc-directed antibody functionalization.

### 2.4. Covalent crosslinking of ADP1 with antibody and retained antigen binding of IgG

Having established that ADP1 successfully guides the non-covalent assembly of intact antibodies, we next sought to transition this system into a covalently crosslinked architecture. We pursued covalent linking for three primary strategic reasons. First, we aimed to covalently lock the ADP1–IgG binding state, thereby improving control over conjugate stoichiometry and limiting time-dependent peptide exchange within the non-covalent network. Second, because the non-covalent assembly is driven by adding two equivalents of ADP1 per full-length antibody, covalent crosslinking allows us to lock the complexed population and efficiently eliminate any residual free ADP1 from the solution. Finally, a covalent bond locks the ADP1–antibody complex to prevent the peptide from releasing, mitigating the risk of peptide dissociation and subsequent off-target exchange with native serum antibodies.

To facilitate this downstream bioconjugation, the ADP1 sequence was deliberately selected from our initial design pool to be entirely devoid of lysine residues. Because this specific sequence lacks primary amines within its framework, it inherently eliminates competing intra-peptide reaction sites. This intentional selection—which served as the singular ADP1 framework across all preceding structural and biophysical characterizations—allowed us to deploy proximity-directed covalent chemistry cleanly, preventing non-specific crosslinking or heterogeneous, off-target aggregation that could disrupt the antibody structure. In the Fc-bound structure, the N-terminus of the parental Z34C domain is positioned in close proximity to Fc Lys246. Given that our X-ray model confirmed that the canonical Z34C-derived binding mode remains strictly conserved in the optimized ADP1 miniprotein, we exploited this geometric arrangement to drive proximity-induced conjugation. Specifically, we employed the short-chain, amine-reactive homobifunctional crosslinker disuccinimidyl glutarate (DSG) to target the primary amine at the unblocked N-terminus of ADP1 for proximity-directed crosslinking to a proximal lysine residue on the Fc surface^6^. Because DSG possesses a short spacer-arm length of 7.7 Å, efficient crosslinking is strongly dependent on spatial proximity, thereby favoring covalent coupling when the peptide is correctly docked within the Fc-binding pocket (**Figure 5A**).

**Figure 5.**
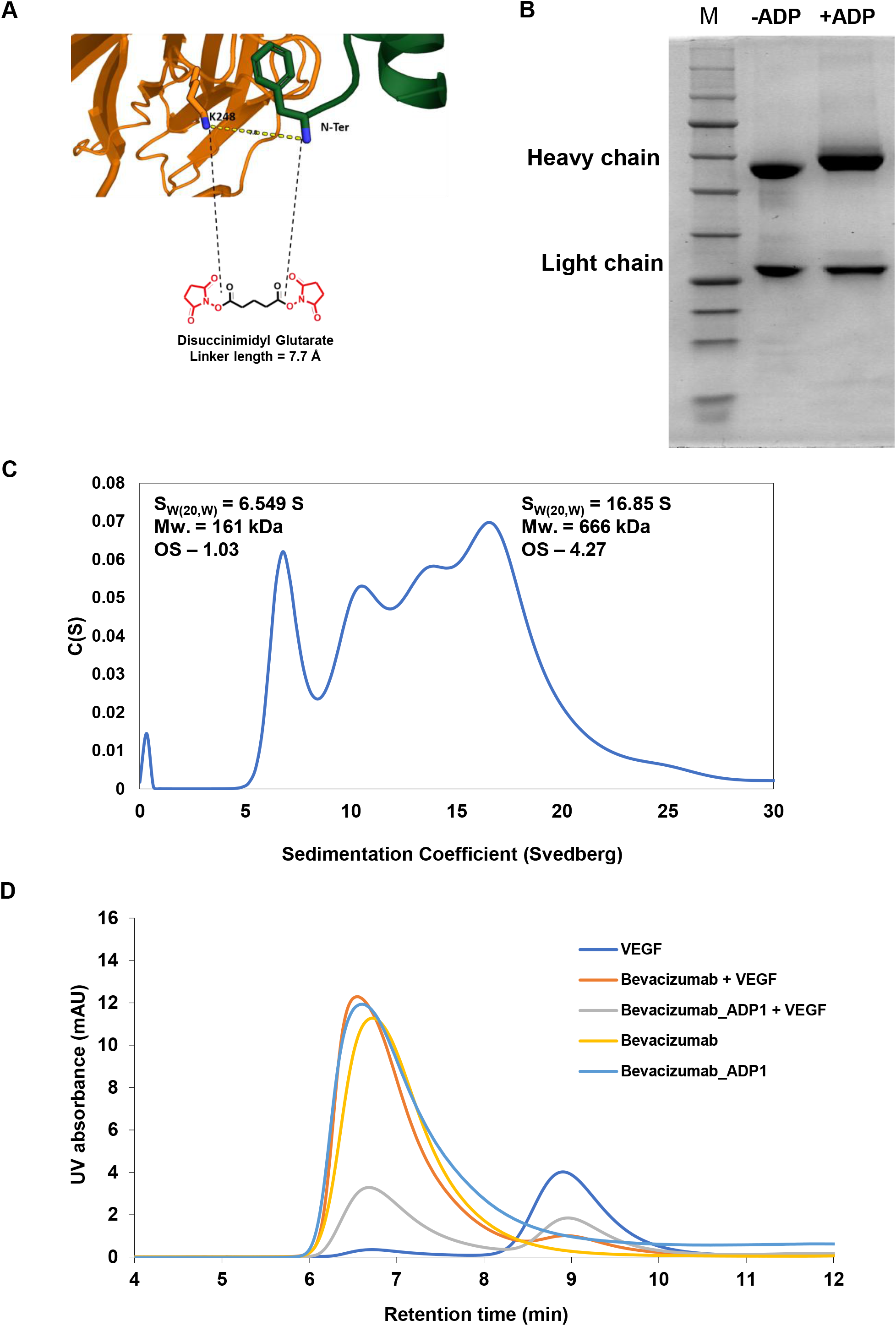
Fc-directed covalent IgG modification by ADP1 and retention of antigen binding. (A) Schematic of Fc-directed covalent ADP1 functionalization. (B) SDS-PAGE analysis of full-length IgG modified with labeled ADP1. (C) Analytical ultracentrifugation analysis of ADP1-conjugated full-length IgG. (D) SEC analysis of ADP1-modified bevacizumab mixed with VEGF.

To evaluate the efficiency and molecular stoichiometry of the crosslinking reaction, we analyzed the products using SDS-PAGE and intact mass spectrometry. While SDS-PAGE revealed a distinct, conjugation-associated mobility shift confined exclusively to the antibody heavy-chain region (**Figure 5B**), intact mass spectrometric analysis of the purified ADP1–IgG conjugate revealed a mass increase consistent with the covalent attachment of two ADP1 protomers per IgG molecule. Conversely, the light chain remained entirely unaltered, demonstrating the excellent site-selectivity of the proximity-guided strategy. Together, these analytical data support predominantly symmetric crosslinking, with one ADP1 module coupled at each of the two Fc-binding sites, without substantial non-specific over-labeling. Following purification, analytical ultracentrifugation (AUC) demonstrated that this covalently locked ADP1–IgG complex maintained its structural capacity for higher-order assembly, populating macromolecular species significantly larger than monomeric IgG (**Figure 5C**).

To assess whether ADP1 modification interfered with antigen recognition, ADP1-modified bevacizumab was mixed with VEGF and analyzed by size-exclusion chromatography. Upon mixing, we observed a substantial decrease in the area under the curve (AUC) for the unbound VEGF peak, matching the depletion profile observed when the antigen was mixed with native, unmodified bevacizumab (**Figure 5D**). This quantitative reduction in the free antigen population indicates that the proximity-induced covalent attachment of ADP1 at the remote Fc Lys246 site proceeds cleanly without altering or sterically occluding the Fab variable domains. These results indicate that ADP1-modified bevacizumab retained detectable VEGF-binding capability under the tested SEC conditions.

Together, these findings extend the ADP1 platform beyond noncovalent antibody assembly. By targeting the Fc region, ADP1 functionalization allows full-length IgG to be modified without directly perturbing the Fab domains, providing a potential route for post-production antibody functionalization and multivalent antibody engineering.

### 2.5. Z34C-mediated modulation of ADP1-driven antibody assembly

Finally, we examined whether the parent Fc-binding peptide Z34C could modulate ADP1-driven antibody assembly. Because ADP1 was derived from Z34C and preserves its Fc-recognition surface, we reasoned that Z34C could compete with ADP1 for overlapping Fc-binding sites on IgG. However, unlike ADP1, Z34C lacks the redesigned peptide–peptide dimerization interface required for Fc-to-Fc propagation. Thus, Z34C was expected to occupy Fc-binding sites without introducing additional peptide-mediated connectivity, thereby acting as a nonpropagating competitor or capper (**Figure 6A**).

**Figure 6.**
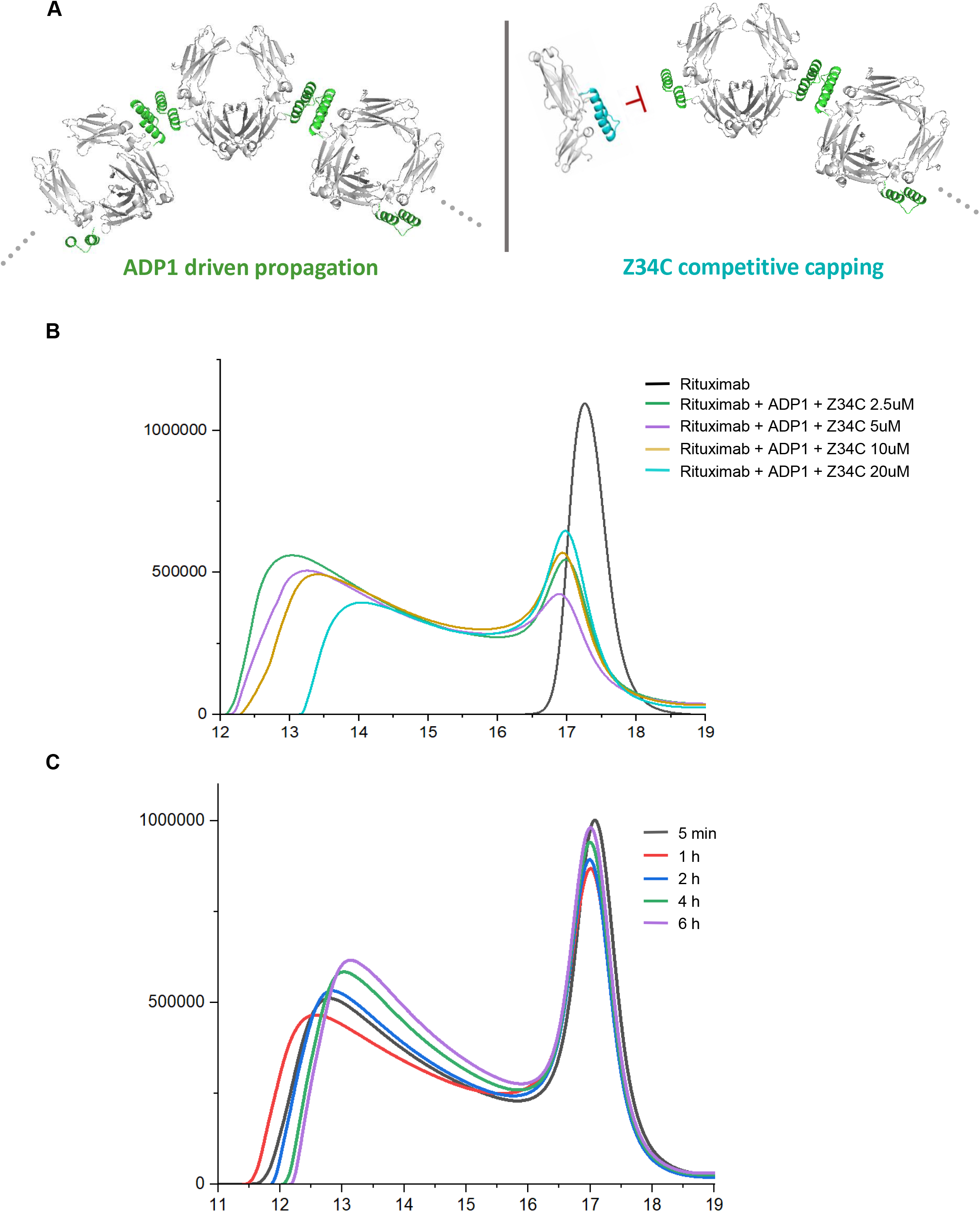
Z34C-mediated modulation of ADP1-driven antibody assembly. **(A)** Schematic of ADP1-driven propagation and Z34C-mediated competitive capping. **(B)** SEC analysis of rituximab incubated with ADP1 in the presence of increasing Z34C concentrations. **(C)** Time-course SEC analysis of the selected rituximab–ADP1–Z34C condition.

To test this concept, rituximab was incubated with ADP1 in the presence of varying Z34C concentrations, and the resulting species distribution was analyzed by size-exclusion chromatography. Under Z34C-containing conditions, the position and intensity of SEC-visible assembly-associated peaks in the 10–16 mL region changed depending on the Z34C concentration (**Figure 6B**). These changes suggest that Z34C influences the ADP1-mediated assembly process and may reshape the assembly distribution through Fc-binding site competition.

We further analyzed a selected Z34C condition by time-course SEC to examine how the ADP1/Z34C-modulated assembly profile evolved over time. Under the selected condition, the early-eluting oligomeric region increased up to 1 h and then gradually shifted toward smaller IgG-containing species at later time points (**Figure 6C**). This time-dependent profile change suggests that ADP1-mediated Fc-to-Fc association may dominate during the early phase, whereas Z34C-mediated Fc-site occupation may progressively reshape the assembly distribution over time.

Together, these results support the concept that Z34C can serve as a secondary modulator of ADP1-driven antibody assembly. By competitively occupying Fc-binding sites without providing a propagation interface, Z34C can alter the distribution of ADP1-containing IgG assemblies. Although SEC profiles alone do not allow precise assignment of the oligomeric state of each peak, these results provide a proof-of-concept strategy for tuning Fc-mediated antibody assembly using the parent Fc-binding peptide.

## 3. Conclusion

In this study, we developed a designed Fc-binding peptide platform that enables Fc-mediated antibody assembly and functionalization without genetic modification of existing IgG molecules. ADP1 was rationally redesigned from the Protein A-derived Fc-binding peptide Z34C as a stable dimeric Fc-binding peptide that preserves the Fc-recognition surface of the parent peptide while introducing peptide–peptide interactions on the opposite helical surface. Through this design, we sought to convert a simple Fc-recognition element into an assembly-driving module that provides connectivity between neighboring Fc regions.

Structural analyses using cryo-EM and X-ray crystallography showed that ADP1 can couple Fc binding with peptide-mediated dimerization to induce spiral higher-order Fc assemblies. The cryo-EM structure demonstrated that ADP1 can organize human serum-derived IgG Fc into an ordered helical assembly, whereas the X-ray structure provided the atomic basis of ADP1–Fc recognition and the ADP1–ADP1 dimerization interface. Although the two structures exhibited distinct helical geometries, both supported the common assembly principle that ADP1 converts Fc recognition into peptide-mediated Fc-to-Fc connectivity.

This assembly principle was further extended to full-length IgG. AUC, SEC, and SEC–MALS analyses provided complementary evidence that ADP1 can promote higher-order antibody association in the context of intact IgG, indicating that the design principle observed in Fc-fragment structures can be translated to full-length antibody systems containing Fab domains. In addition, Fc-directed covalent ADP1 functionalization enabled modification of full-length IgG while retaining VEGF binding by bevacizumab, demonstrating that ADP1 can serve not only as an antibody assembly module but also as a functionalization handle.

The parent Z34C peptide was also explored as a secondary modulator of ADP1-driven antibody assembly. Under Z34C-containing conditions, SEC-visible assembly-associated peaks changed depending on Z34C concentration and incubation time, suggesting that competitive Fc-site occupation can modulate the ADP1-mediated assembly distribution. However, because SEC profiles alone do not allow precise assignment of the oligomeric state of each peak, these results should be interpreted as a proof-of-concept for Z34C-mediated modulation.

Overall, this work shows that rational redesign of an Fc-binding peptide can extend Fc recognition into multivalent antibody assembly and Fc-directed functionalization. The ADP1-based approach provides a modular post-production strategy for organizing and functionalizing existing antibodies without genetic engineering of the IgG scaffold. In future studies, ADP1-mediated Fc association may be explored in applications where multivalent antibody presentation is advantageous, such as diagnostic signal amplification or antibody engineering. In addition, the ability of ADP1 to induce Fc-mediated IgG association may be further explored as a potential strategy for selective IgG enrichment or isolation from complex biological fluids such as serum. However, such applications will require further validation of assembly reversibility, IgG selectivity, serum compatibility, and recovery and purity in complex samples.

## 4. Methodology

### 4.1. Computational design of ADP1

ADP1 was computationally designed using the reported complex structure of the Protein A-derived Z34C peptide bound to the IgG Fc region (PDB ID: 5U52) as the starting scaffold. The Z34C peptide was extracted from the complex and used as the parent Fc-binding peptide. To generate an assembly-driving Fc binder, two Z34C peptide chains were manually positioned to create an initial dimeric architecture in which the Fc-binding faces remained compatible with Fc recognition and the opposite helical surfaces were oriented toward each other for peptide–peptide interface design.

ProteinMPNN was used to redesign the peptide dimer interface while fixing residues involved in Fc interaction to preserve the parent Fc-binding mode. Sequence outputs were evaluated using AlphaFold-Multimer to assess maintenance of the intended dimeric architecture and compatibility with Fc binding. An additional symmetric design round was then performed using tied residue constraints at the designed interface positions, while Fc-interacting residues were fixed in both peptide chains. Candidate sequences were prioritized based on predicted maintenance of Fc-binding geometry, formation of the intended dimer interface, overall structural quality, and predicted solubility.

Top candidate models were further inspected, and one high-priority design was selected for additional optimization. Point substitutions at the terminal cysteine position were evaluated to reduce unfavorable terminal interactions and improve the predicted dimer interface. Among the tested variants, C33V was selected for the final design. The resulting 34-residue peptide was designated ADP1 and used for experimental characterization. Detailed fixed and tied residue positions used for ProteinMPNN design are provided in **Supplementary Table 1**.

### 4.2. Peptide synthesis and purification

Peptides were synthesized on Rink Amide AM resin (0.7 mmol g^−1^ loading capacity) on a 150 mg scale using a standard Fmoc-based solid-phase peptide synthesis (SPPS) strategy as described previously^12^. Global deprotection and cleavage of the resin-bound peptides were carried out in a cleavage cocktail consisting of 90% trifluoroacetic acid (TFA), 7% triisopropylsilane (TIS), and 3% water for 2 h at room temperature. The deprotected crude peptide was precipitated by the addition of ice-cold diethyl ether. The resulting white solid was pelleted by centrifugation, the supernatant was discarded, and the crude pellet was dissolved in a solvent mixture of 25% dimethyl sulfoxide (DMSO), 25% acetonitrile (ACN), and 50% water^13^. To initiate disulfide bond formation, the solution was adjusted to pH 7.0 by the addition of 500 mM Tris buffer and incubated overnight at room temperature to complete cyclization.

Purifications were performed using a semipreparative column (Phenomenex C18, 250 × 10 mm, 110 Å, 5 µm) at a flow rate of 4 ml min^−1^ by reversed-phase–high-performance liquid chromatography (RP–HPLC) using a gradient of 0.1% TFA in water and 0.1% TFA in acetonitrile. The purity was checked using an analytical column (Phenomenex C18, 250 × 4.6 mm, 100 Å, 5 µm) at a flow rate of 1 ml min^−1^ by RP–HPLC and the identity was confirmed by electrospray ionization–mass spectrometry.

### 4.3. Circular dichroism spectroscopy

Far-UV circular dichroism spectra of the peptides were recorded on a JASCO-815 spectropolarimeter using a 0.1 cm path-length cuvette at 50 μM concentration in 20 mM sodium-phosphate buffer (pH 7.4). Scans were carried out at 20 and 95 °C and again at 20 °C after fast refolding (10 min) over the range of 190–260 nm with 0.5-nm increments and a 2-nm bandwidth. Thermal unfolding experiments were monitored at 222 nm over a temperature range of 4–96 °C at 1 °C interval in 20 mM sodium-phosphate buffer (pH 7.4). The samples were equilibrated for 30 seconds at each temperature. Temperature-dependent circular dichroism data were fit to a two-state unfolding model to obtain the melting temperature (*T*_m_).

### 4.4. Reduction kinetics

To evaluate the chemical stability of the engineered disulfide architecture, the reduction kinetics of the cyclic peptides were monitored directly via automated chromatographic analysis. The cyclic peptide was incubated in a solution containing 2 mM dithiothreitol (DTT) at room temperature (25 ^∘^C). Reaction progress was tracked by direct, unquenched injections onto the high-performance liquid chromatography (HPLC) system at specified time intervals, beginning at 5 min post-initiation and proceeding at fixed 35 min intervals thereafter. Aliquots were resolved using reversed-phase HPLC (RP-HPLC) on a C18 column, where the acidic mobile phase served to arrest further reduction upon injection. The relative abundance of the intact cyclic peptide at each time point was quantified by integrating the peak area under the curve corresponding to the cyclic species. The resulting values were plotted as a function of incubation time and fitted to a non-linear one-phase exponential decay model to calculate the kinetic half-life (t_1/2_) of the reduction process.

### 4.5. Proteolytic stability assay

Peptides (50 µM in Tris buffered saline, pH 7.4) were incubated with proteinase K (10 µg/ml final concentration) at room temperature. The reaction at each time point was manually quenched at the addition of 50 ul of reaction mixture to 200ul of 1% TFA in water. The quenched samples were analysed using analytical RP-HPLC on a C18 column. The area under the curve of the starting peptide peak monitored at 220 nm was calculated for each time point, plotted against incubation time, and fitted to a one-phase exponential decay model to determine the t ^14^.

### 4.6. Preparation of human IgG Fc by papain digestion and purification

Papain was first activated by incubation in an activation buffer (10mM Tris, 10mM L-cysteine hydrochloride, 2 mM EDTA, pH 7.2) for 2 h at room temperature with agitation at 300 rpm. To perform the subsequent enzymatic cleavage under non-reducing conditions, the activated papain was buffer-exchanged into a cysteine-free digestion buffer (100mM Tris, 2 mM EDTA, pH 7.2)^9,10^. Intact human IgG (5mg/ml) was then incubated with the activated papain at a 50: 1 substrate-to-enzyme weight ratio for 4 h at 37^∘^C and 300 rpm. Following complete digestion, the papain enzyme (∼ 23 kDa) was separated from the reaction mixture by centrifugal ultrafiltration using a 30 kDa molecular weight cut-off (MWCO) filter. The resulting retentate was collected, yielding a mixture containing the cleaved Fc (∼ 50 kDa) and Fab (∼ 50 kDa) fragment.

To isolate the cleaved Fc fragment from the digestion mixture, affinity chromatography was performed using immobilized Protein A agarose beads (786-283, G-Biosciences) following the manufacturer’s prescribed protocol. As a final polishing step to ensure high conformational homogeneity and remove trace aggregates, residual uncleaved antibody, or over-cleaved Fc contaminants, size-exclusion chromatography (SEC) of the isolated Fc was performed. The neutralized Protein A eluate was loaded onto a Superdex 75 (S75) column pre-equilibrated and eluted with 1x PBS buffer (pH 7.4). Eluted fractions corresponding to the monomeric human IgG Fc peak were pooled and quantified via UV absorbance at 280 nm. Purified Fc fragments were stored at −80^∘^C for short-term experimental use or lyophilized for long-term storage.

### 4.7. ITC binding analysis of peptides with human IgG Fc

Thermodynamic binding profiles between the synthesized peptides and human IgG Fc fragments were determined via isothermal titration calorimetry (ITC) using a MicroCal calorimeter at 25^∘^C. Both the peptide and human IgG Fc samples were prepared in 1 × PBS buffer (pH 7.4). Titrations were performed via an initial injection of 0.4 μL followed by 18 subsequent automated injections of 2 *μ*L spaced at regular intervals of 150 seconds, with continuous stirring at 750 rpm. Heats of dilution obtained from peptide titration into buffer were subtracted from the corresponding peptide-into-Fc titration data. The net integration values of the raw thermograms were plotted as heat per injection against the molar ratio and fitted to an independent binding model using the instrument’s analysis software to calculate the dissociation constant (*K_d_*), stoichiometry (*n*), and thermodynamic binding parameters (Δ*H* and Δ*S*).

### 4.8. Analytical ultracentrifugation

The sedimentation-velocity analytical ultracentrifugation data for all samples were adjusted to an absorbance of 0.3–0.9 at 280 nm in a 10 mm path-length cell using an Optima XL-I analytical ultracentrifuge equipped with absorbance optics with an An-60Ti 4 place rotor (Beckman Inc.). Peptides were dissolved in the 20 mM Phosphate buffer pH 7.4 and the antibody containing sample were dissolved in 20 mM Phosphate buffer with 150mM NaCl pH 7.4. Sedimentation-velocity experiments were carried out at 30000 or 40,000 rpm at 20 °C using two-channel charcoal-filled centerpieces with Sapphire glass windows. Samples were loaded onto the two-sector centerpiece (400 µl of reference cells and 390 µl of sample cells). The velocity data were collected by scanning samples at a wavelength of 280 nm with a spacing of 0.003 cm and an average of three scans per step. The standard partial specific volumes of peptides and proteins (roughly 0.73 cm^3^ g^−1^ at 20 °C) and buffer density (roughly 1.00 g cm^−3^ at 20 °C) and viscosity (roughly 0.01002 poise at 20 °C) were calculated using the program SEDNTERP. Oligomeric states of peptides and complexes were analyzed by direct curve fitting of sedimentation boundaries using Sedfit^15^. Fit to data was selected based on the root mean square deviations (rmsd) less than 0.008. The sedimentation coefficients were normalized to 20 °C in water, *S*_20,w_ under standard conditions.

### 4.9. Fluorescence-detection Size-exclusion chromatography analysis of Fc and IgG assemblies

To monitor the formation and structural distribution of higher-order macromolecular complexes, analytical fluorescence size-exclusion chromatography (FSEC) was performed. Human IgG Fc fragments (10 *μ*M) or full-length IgG molecules (3 *μ*M) were mixed with the peptide conjugate at a 1: 2 protein-to-peptide molar ratio in 1 × PBS buffer (pH 7.4). The reaction mixtures were incubated at room temperature for 30 min. Following incubation, the samples were resolved on a 3 mL Superdex 200 (S200) analytical column pre-equilibrated and eluted with 1 × PBS buffer. Elution profiles were tracked via real-time FSEC monitoring of native tryptophan fluorescence (excitation at 290 nm, emission at 334 nm) to visualize the formation of macromolecular assemblies through the shift in retention volume of the antibody species. Since the Z34C and ADP1 peptide sequences lack tryptophan residues, excess or unbound peptide variants remain fluorescence-invisible under these optical parameters, ensuring that the monitored signal corresponds exclusively to the protein-containing species and their higher-order complexes.

### 4.10. SEC–MALS analysis

Size-exclusion chromatography coupled with multi-angle light scattering (SEC-MALS) was used to determine the molar masses and oligomeric distributions of Fc–ADP1 and full-length IgG–ADP1 assemblies. Three sample types were analyzed: commercial human serum-derived IgG Fc complexed with ADP1, in-house-prepared bevacizumab Fc complexed with ADP1, and full-length bevacizumab complexed with ADP1. For all three sample types, the Fc fragment or full-length bevacizumab was mixed with ADP1 at a 1:2 protein-to-peptide molar ratio. The final concentrations of the Fc fragment or full-length bevacizumab and ADP1 were 56 and 112 μM, respectively. Samples were prepared in 10 mM sodium phosphate buffer containing 50 mM NaCl at pH 7.4. SEC-MALS analysis was performed at 25°C using an Agilent 1260 Infinity II HPLC system coupled to a Wyatt DAWN multi-angle light-scattering detector and a differential refractive-index detector. Samples (500 μL) were injected onto a TSKgel G4000 column with a nominal separation range of 20–7,000 kDa, pre-equilibrated and eluted with the same buffer at a flow rate of 0.45 mL min⁻¹. The column eluate was monitored by multi-angle light scattering and differential refractive-index detection. Molar masses across the elution peaks were calculated from the combined light-scattering and concentration signals using ASTRA software and the standard protein refractive-index increment, dn/dc, of 0.185 mL g⁻¹.

### 4.11. Cryo-EM sample preparation, data collection, and image processing

For cryo-EM sample preparation, commercial human serum-derived IgG Fc was incubated with ADP1, and the resulting mixture was subjected to size-exclusion chromatography to isolate the higher-order Fc–ADP1 assembly. Fractions corresponding to the early-eluting Fc–ADP1 complex peak were collected and pooled **(Supplementary Figure 3)**. The pooled sample was concentrated to approximately 1 mg mL⁻¹ using a centrifugal concentrator with a 100 kDa molecular-weight cutoff.

The purified Fc–ADP1 complex was applied to glow-discharged holey-carbon copper grids (Quantifoil R1.2/1.3, 200 mesh; EMS). The grids were blotted for 3.0 s and plunge-frozen in liquid ethane cooled with liquid nitrogen using a Vitrobot Mark IV (Thermo Fisher Scientific), with the environmental chamber maintained at 100% humidity and 4°C. Automated cryo-EM data acquisition was performed at the Korea Research Institute of Bioscience and Biotechnology (KRIBB), Daejeon, Republic of Korea, using Glacios transmission electron microscope (Thermo Fisher Scientific) operated at an accelerating voltage of 200 kV. Data acquisition was performed using EPU software (Thermo Fisher Scientific), and images were recorded using a Falcon IV direct electron detector (Thermo Fisher Scientific). Total 4,877 micrographs were collected at a defocus range of −1.4 to −2.2 μm, at a nominal magnification of 92,000×, corresponding to a physical pixel size of 1.1 Å. Each exposure was dose-fractionated into 40 movie frames over a total exposure time of 8.18 s, resulting in a total electron dose of 39.8 e⁻ Å⁻².

Cryo-EM data processing was performed using cryoSPARC version 4.7.1. Movie frames were aligned and dose-weighted to correct for beam-induced motion, and contrast transfer function parameters were estimated from the motion-corrected micrographs. Particles were initially selected manually and subjected to two-dimensional classification. Representative class averages were used as templates for filament tracing, and the resulting particles were further classified in 2D. Well-defined 2D class averages were subsequently used for template-based automated particle picking, yielding 4,715,554 extracted particle images. After 2D classification, the selected particles were subjected to ab initio reconstruction and homogeneous refinement, followed by final helical refinement. The final Fc–ADP1 reconstruction was obtained at an overall resolution of 4.88 Å, as estimated using the gold-standard Fourier shell correlation criterion at FSC = 0.143.

A structural model of the Fc–ADP1 complex was fitted into the cryo-EM density map using UCSF ChimeraX version 1.10. The fitted model was manually adjusted in WinCoot and subjected to restrained real-space refinement using phenix.real_space_refine in PHENIX version 1.21.2-5419. Within the higher-order assembly, the Fc–ADP1 units adopted an elongated and distorted conformation relative to the reference structure, precluding reliable refinement of a complete atomic model. The fitted model was therefore used only to facilitate interpretation of the overall assembly architecture and was not considered a fully refined atomic model. Accordingly, only the cryo-EM density map and associated half-maps were deposited.

### 4.12. Crystallization, X-ray data collection, and structure determination

Bevacizumab-derived Fc and ADP1 were mixed at final concentrations of 115 and 230 μM, respectively, corresponding to a 1:2 Fc-to-peptide molar ratio. The mixture was incubated for 1 h at room temperature and subsequently concentrated to 5 mg mL⁻¹ in 10 mM sodium phosphate buffer containing 50 mM NaCl. Crystallization screening was performed using a Mosquito liquid-handling robot by the sitting-drop vapor-diffusion method.

Crystals suitable for X-ray diffraction were obtained in a reservoir solution containing 1260 mM Ammonium sulfate, 100 mM Sodium acetate/Acetic acid pH 4.5, 200 mM Sodium Chloride. Prior to X-ray data collection, crystals were briefly soaked in a cryoprotectant solution consisting of the reservoir solution supplemented with 50% (v/v) glycerol, and then flash-cooled directly in liquid nitrogen.

X-ray diffraction data were collected to a resolution of 2.60 Å. The crystals belonged to space group P6₁, with unit-cell parameters of a = b = 117.64 Å, c = 127.21 Å, α = β = 90°, and γ = 120°. Diffraction data were indexed, integrated, and scaled using the data-processing software employed at the beamline. Data collection and refinement statistics are summarized in **Supplementary Table 4**.

The structure was solved by molecular replacement using a predicted 3D structure generated by AlphaFold3 as the search model. Iterative model building and manual correction were performed using WinCoot, and refinement was carried out using PHENIX version 1.20.1-4487. Successive rounds of model refinement and manual inspection were performed until convergence. The final structure was refined to R_work and R_free values of 0.219 and 0.264, respectively. Model geometry and stereochemical quality were evaluated using the validation tools implemented in PHENIX. The final coordinates and structure factors were deposited in the Protein Data Bank under accession code 27UM.

### 4.13. Covalent conjugation of ADP1 to IgG

The covalent modification of ADP1 was adapted from previously described literature methods^6^. A 500 *μ*L aliquot of the peptide (2 mM in DMSO containing 2% pyridine) was mixed with 500 *μ*L of disuccinimidyl glutarate (DSG, 50 mM in acetonitrile). The reaction mixture was incubated at 50^∘^C for 3 h to yield the functionalized succinimidyl glutararyl (SG)-peptide derivative. The reaction was quenched by diluting it 5-fold in 5% trifluoroacetic acid (TFA) in acetonitrile and purification was performed using a semipreparative column (Phenomenex C18, 250 × 10 mm, 110 Å, 5 µm) at a flow rate of 4 ml min^−1^ by reversed-phase–high-performance liquid chromatography (RP–HPLC) using a gradient of 0.1% TFA in water and 0.1% TFA in acetonitrile. The fractionated product peaks were collected and immediately lyophilized to a dry powder. The resulting purified powder was immediately dissolved in 0.1% TFA in DMSO and stored at −80^∘^C until further use.

For the antibody-peptide conjugation step, the full-length IgG (5 *μ*M) was dissolved in a conjugation buffer consisting of 20 mM sodium phosphate, 500 mM NaCl, and 5% glycerol (pH 7.0). The coupling reaction was initiated by adding 3 equivalents of the activated, labelled peptide and incubating the mixture at 20^∘^C for 8 h under continuous agitation at 300 rpm. To drive the coupling reaction to completion while mitigating hydrolysis of the succinimidyl glutarate group on the peptide, an additional 1 equivalent of the activated peptide was added to the reaction mixture, followed by a final overnight incubation at 4^∘^C.

### 4.14. SEC analysis of VEGF binding by ADP1-modified bevacizumab

To evaluate the functional antigen-binding capability of the bevacizumab–ADP1 covalent conjugate, analytical size-exclusion chromatography (SEC) was performed. Unmodified bevacizumab or the bevacizumab–ADP1 covalent conjugate ( 0.75 *μ*M) was mixed with vascular endothelial growth factor (VEGF, 1.5 *μ*M) in 1 × PBS buffer (pH 7.4) and incubated at room temperature for 30 min to allow complex formation to reach equilibrium. Following incubation, the reaction mixtures were resolved on a 3 mL Superdex 75 (S75) analytical column pre-equilibrated and eluted with 1 × PBS buffer. Elution profiles were monitored via continuous UV absorbance at 220 nm. Under these chromatographic conditions, all high-molecular-weight antibody species—including free IgG, the bevacizumab–ADP1 covalent conjugate, and their respective VEGF immunocomplexes—exceed the exclusion limit of the resin (∼ 70 kDa) and elute cleanly within the column void volume (*V*_0_). Unbound, free VEGF (∼ 24 kDa) elutes well within the linear resolving range of the column. This separation enabled VEGF sequestration to be evaluated by measuring depletion of the free VEGF peak relative to control samples.

### 4.15. SEC analysis of Z34C-mediated modulation of ADP1-driven antibody assembly

To evaluate the effect of Z34C on ADP1-mediated antibody assembly, ADP1 and Z34C were first mixed at final concentrations of 20 μM and 1.25, 2.5, or 10 μM, respectively. Rituximab was then added to a final concentration of 10 μM, corresponding to a 1:2 antibody-to-ADP1 molar ratio. The resulting mixtures were incubated for 1 h at room temperature and subsequently analyzed by analytical size-exclusion chromatography.

For time-course analysis, rituximab, ADP1, and Z34C were mixed at final concentrations of 40, 80, and 5 μM, respectively. The mixtures were incubated at room temperature and analyzed after 5 min and 1, 2, 4, and 6 h. All samples were analyzed using the same SEC column, buffer composition, flow rate, injection volume, and UV-detection conditions described above.

## Author Information

### Author Contributions

**Subhrodeep Saha**: Writing – review & editing, Validation, Software, Investigation, Formal analysis, Data curation, Visualization. **Su-Jin Lee**: Writing – original draft, Writing – review & editing, Software, Investigation, Formal analysis, Data curation, Visualization. **Hyun-Jo Shim**: Visualization, Investigation, Data curation. **Song-Yi Jang**: Investigation, Data curation. **Kwang-Hyun Park**: Software, Investigation, Formal analysis. **Ho-Chul Shin**: Software, Investigation, Formal analysis, Supervision. **Eui-Jeon Woo:** Writing – review & editing, Supervision, Project administration, Funding acquisition, Conceptualization. **Jayanta Chatterjee:** Writing – review & editing, Supervision, Project administration, Funding acquisition, Conceptualization. All authors have given approval to the final version of the manuscript. S-J.L. and S.S. contributed equally.

### Notes

A patent application related to the ADP1-based Fc-binding peptide platform for Fc-mediated antibody assembly and functionalization described in this manuscript has been filed by the authors and/or their institutions.

## Acknowledgments

J.C. acknowledges funding from the Indian Institute of Science (IISc) and the DBT–IISc Partnership Program. The authors also acknowledge infrastructural support from DST-FIST, UGC-CAS, the DBT–IISc Partnership Program, and the Ministry of Human Resource Development (MHRD). This work was supported by National Research Foundation of Korea (NRF) grants funded by the Korean Government (MSIP) (RS-2022-NR071772, RS-2021-NR059435, and RS-2021-NR056526 to W.-E.J.), the National Research Council of Science & Technology (NST) (CRC22024-500 to W.-E.J. and S.-H.C.), and the KRIBB Research Initiative Program (KGM5382632, KGM1062612, and KGM1322612 to W.-E.J.). The use of the cryo-EM facilities of the NEXUS consortium was supported by an NRF grant (RS-2024-00440289).

## Supporting information

**Supplementary Table 1.** Design constraints used for ADP1 generation.

| Design step | Chain(s) | Constraint type | Residue positions | Purpose |
| --- | --- | --- | --- | --- |
| Initial ProteinMPNN design | A, B | Fixed | Original numbering:<br>6, 7, 8, 10, 11, 12, 14,<br>15, 16, 18, 19, 25, 28,<br>29, 32, 35, 36, 39 | Preserve Fc-interacting residues from the parent Z34C peptide |
|  |  |  | Renumbered 34-residue peptide: 1, 2, 3, 5, 6, 7,<br>9, 10, 11, 13, 14, 20, 23,<br>24, 27, 30, 31, 34 |  |
| Symmetric ProteinMPNN design | A, B | Fixed | 1, 2, 3, 5, 6, 7, 9, 10, 11,<br>13, 14, 20, 23, 24, 27,<br>30, 31, 34 | Preserve Fc-binding mode during symmetric interface redesign |
|  |  | Tied | 4, 8, 12, 15, 16, 17, 18,<br>19, 21, 22, 25, 26, 28,<br>29, 32, 33 | Enforce sequence symmetry across the designed homodimer interface |

**Supplementary Table 2.** Thermal stability, helicity, and Fc-binding affinity of Z34C, ADP1, and ADP1-derived variants.

| Peptide | $T_m$ (°C) | Helicity (%) | $K_D$, Fc binding (nM) |
| --- | --- | --- | --- |
| Z34C | 59.4 | 49 | 70.8 |
| ADP1 | >95 | 41 | 92.1 |
| ADP1-LA | 64.7 | 54 | 64.6 |
| ADP1-IA | >95 | 42 | - |

**Supplementary Table 3.**
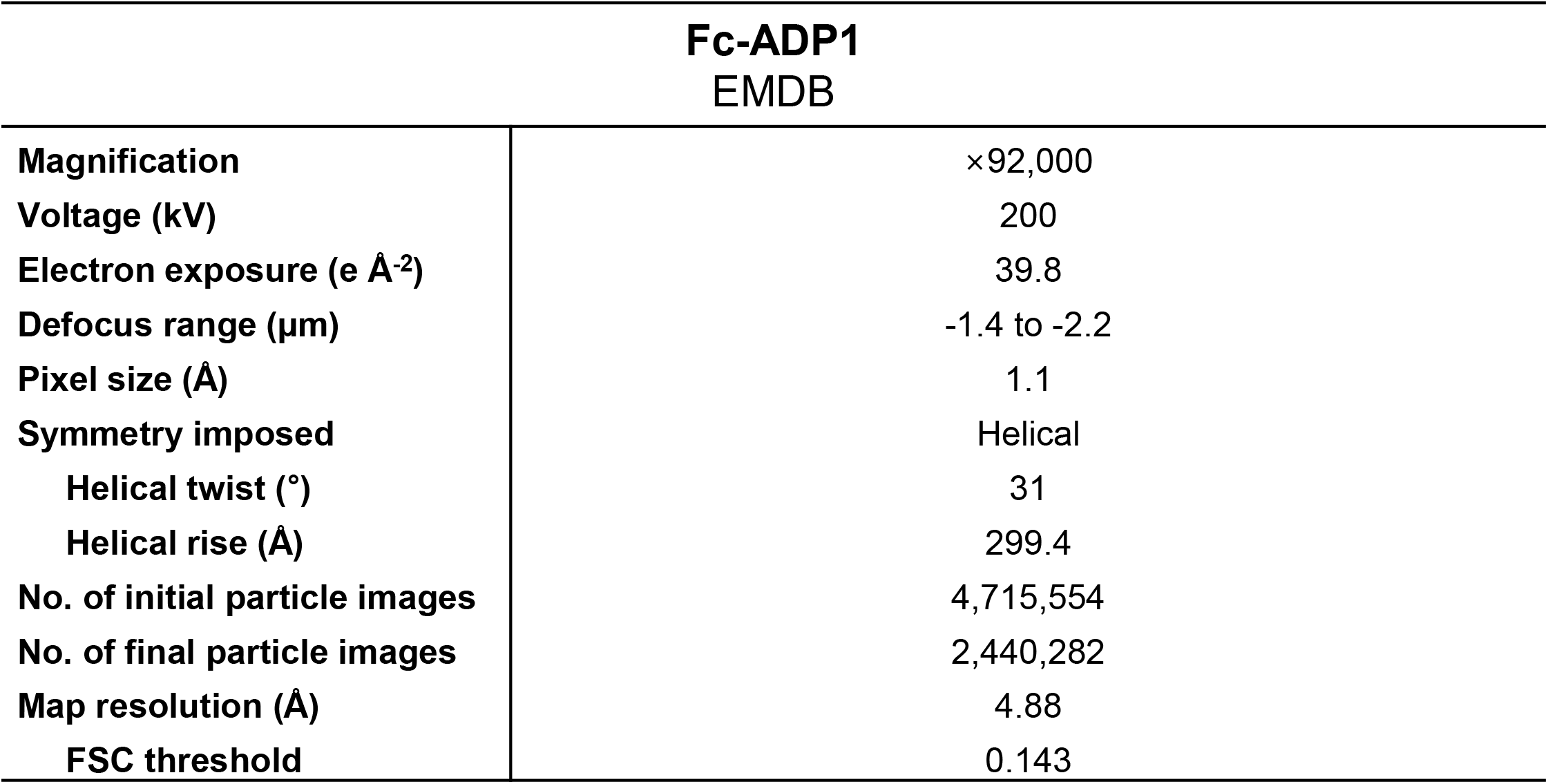
Cryo-EM data collection statistics.

**Supplementary Table 4.** X-ray data collection and refinement statistics.

| ADP1-bevacizumab Fc complex<br>(PDB: 27UM) |  |
| --- | --- |
| Resolution range (Å) | 29.41 - 2.60 (2.68 - 2.60) |
| Space group | P6<sub>1</sub> |
| Unit cell | a = 117.64 Å, b = 117.64 Å, c = 127.21 Å, α = 90°, β = 90°, γ = 120° |
| Unique reflections | 29,906 (2,310) |
| Multiplicity | 20.4 (21.8) |
| Completeness (%) | 97.2 (100.0) |
| Mean I/sigma(I) | 21.57 (5.13) |
| Wilson B factor | 58.2 |
| R-merge | 0.128 (0.811) |
| R-meas | 0.132 (0.830) |
| CC1/2 | 0.999 (0.858) |
| Reflections used in refinement | 29,877 (2,801) |
| Reflections used for R-free | 1,495 (140) |
| R-work | 0.219 (0.316) |
| R-free | 0.264 (0.345) |
| Number of non-hydrogen atoms | 4,447 |
| Macromolecules | 3,916 |
| Glycans/ligands/ions |  |
| Solvent | 283 |
| Protein residues | 483 |
| RMS(bonds) | 0.003 Å |
| RMS(angles) | 0.486° |
| Ramachandran favored (%) | 98.53 |
| Ramachandran allowed (%) | 1.47 |
| Ramachandran outliers (%) | 0 |
| Rotamer outliers (%) | 0 |
| Clash score | 3.29 |
| Average B-factor (Å<sup>2</sup>) | 66.33 |
| Macromolecules | 66.41 |
| Solvent | 45.07 |

Statistics for the highest-resolution shell are shown in parentheses.

---

**Supplementary Figure 1.**
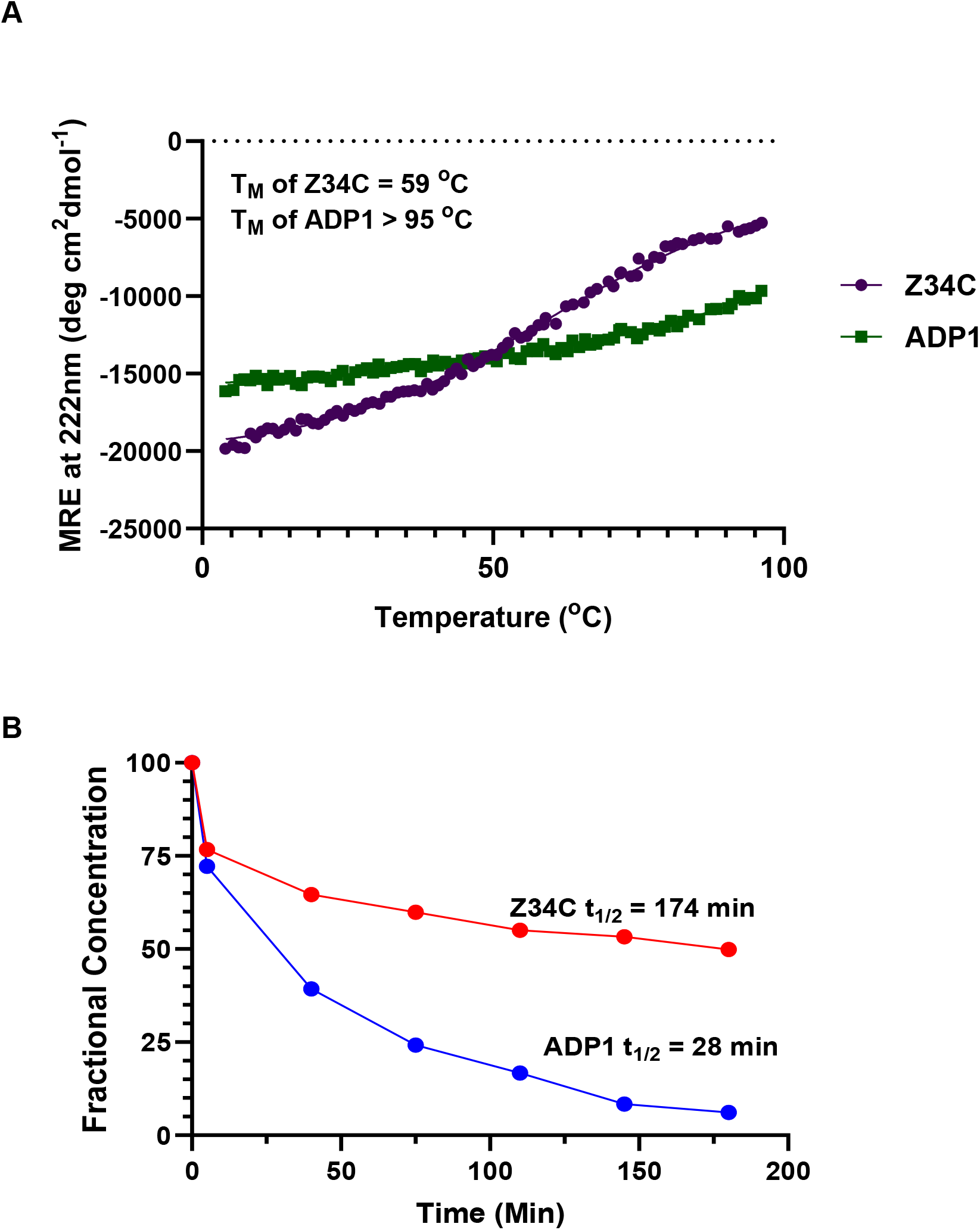
Stability and secondary-structure characterization of ADP1. **(A)** Thermal denaturation of Z34C and ADP1 **(B)** Autocyclization of Z34C and ADP1

**Supplementary Figure 2.**
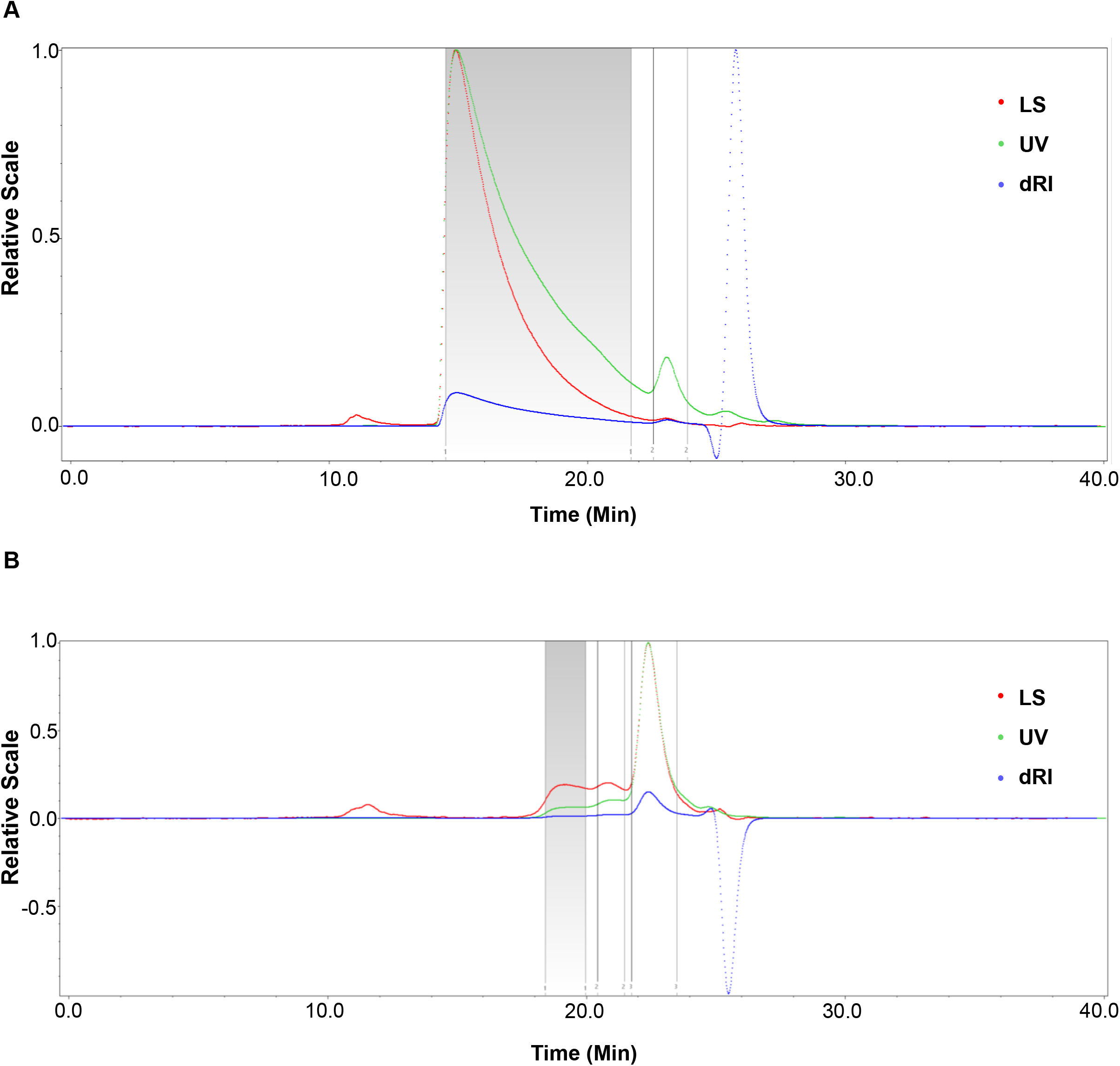
SEC characterization of ADP1-induced Fc assembly. **(A)** SEC-MALS of human serum derived Fc complexed with ADP1 **(B)** SEC-MALS of Bevacizumab Fc complexed with ADP1

**Supplementary Figure 3.**
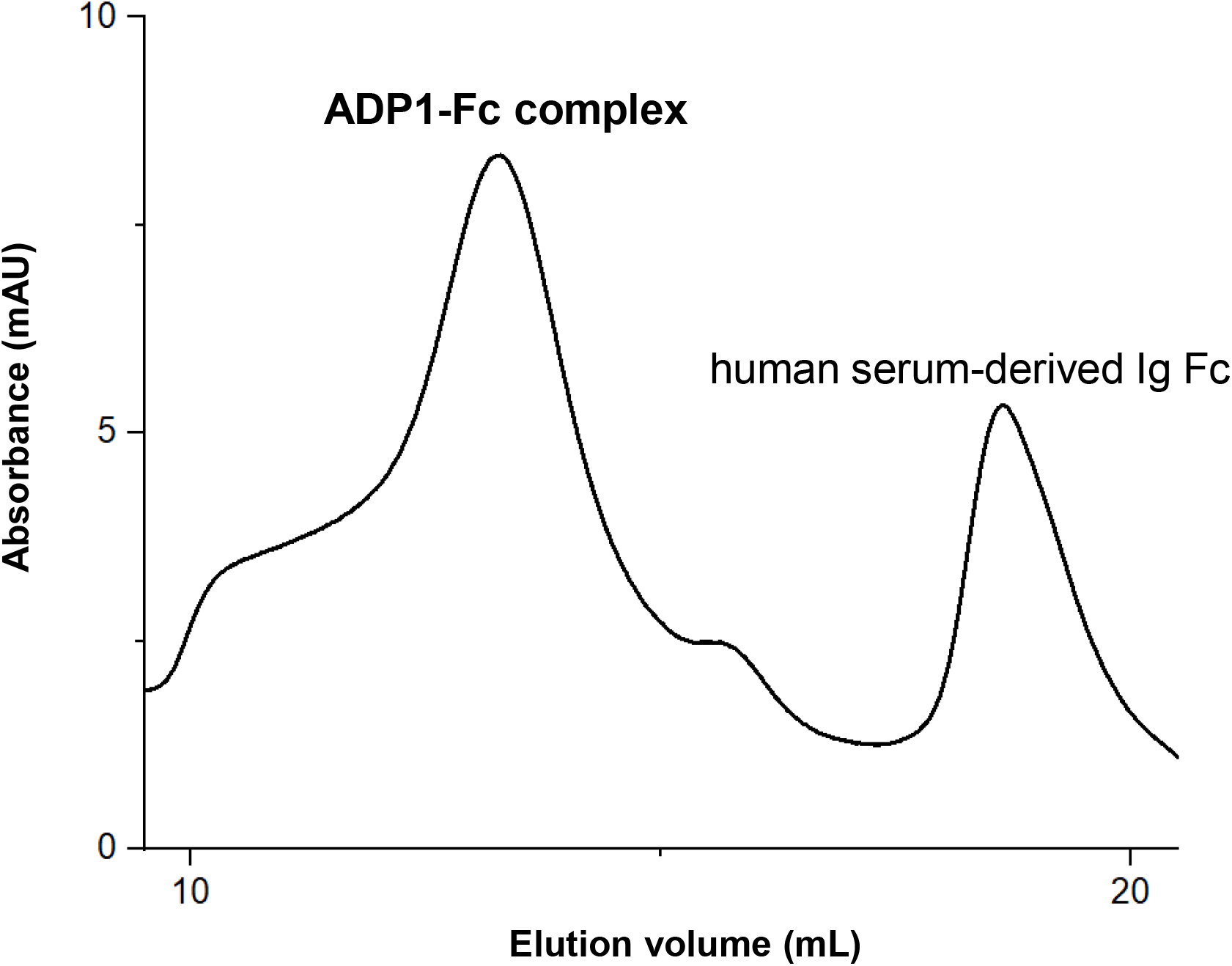
SEC characterization of ADP1-induced Fc assembly. Size-exclusion chromatography analysis of human serum-derived IgG Fc incubated with ADP1.

**Supplementary Figure 4.**
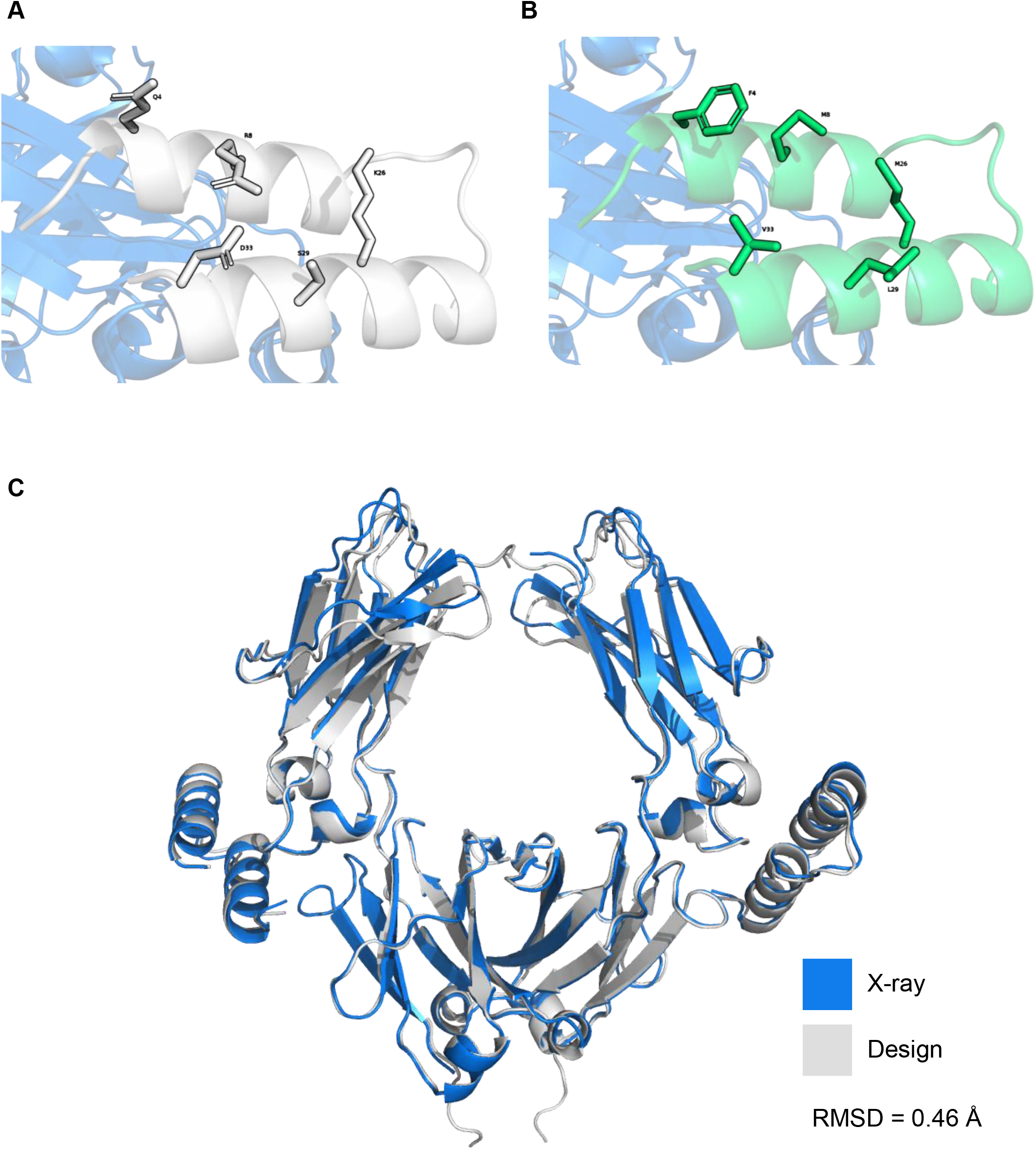
Structural basis of Z34C-to-ADP1 redesign. **(A)** Parent Z34C–Fc structure showing residues selected for redesign on the non-Fc-binding helical surface. Fc is shown in blue and Z34C in gray. **(B)** Redesigned ADP1–Fc structure showing hydrophilic-to-hydrophobic substitutions on the non-Fc-binding peptide surface. Fc is shown in blue and ADP1 in green. **(C)** Overlay of the designed ADP1–Fc model and the experimentally determined ADP1–Fc X-ray structure. X-ray structure, blue; design model, gray. RMSD = 0.46 Å.

**Supplementary Figure 5.**
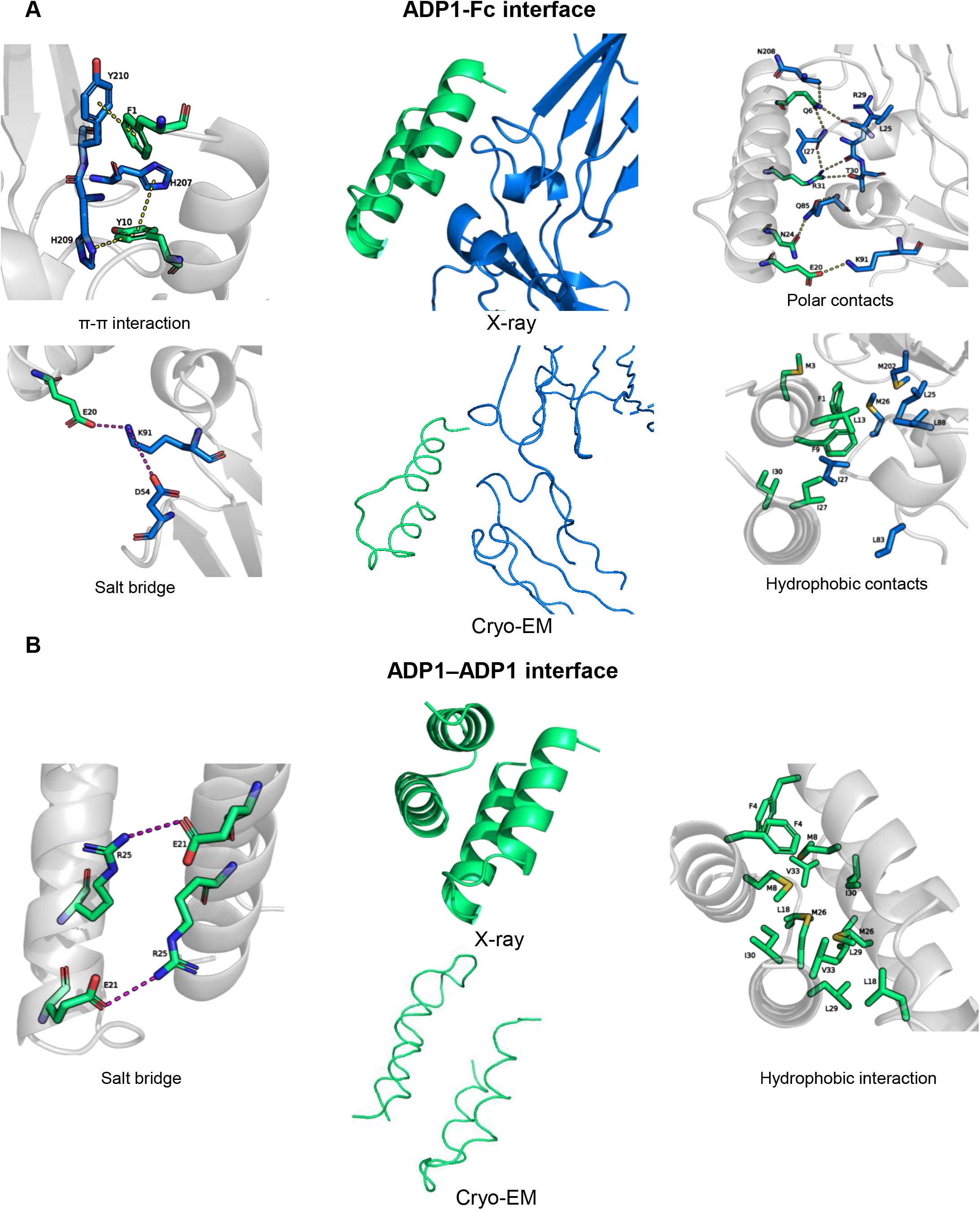
Interface analysis of the ADP1–Fc and ADP1–ADP1 contacts. **(A)** ADP1–Fc interface analysis based primarily on the ADP1–bevacizumab Fc X-ray structure, with the corresponding cryo-EM model shown for comparison. Insets show representative π–π interactions, polar contacts, salt bridges, and hydrophobic contacts. **(B)** ADP1–ADP1 dimerization interface analysis based on the X-ray structure, with the corresponding cryo-EM arrangement shown for comparison. Insets show representative salt-bridge and hydrophobic interactions. Dashed lines indicate polar or electrostatic interactions.

## Notes

### Summary of Updates

The name of co-author Song Yee Jang has been corrected from Song-Yi Jang to Song Yee Jang.

## References

(1) Ku, Z.; Xie, X.; Hinton, P. R.; Liu, X.; Ye, X.; Muruato, A. E.; Ng, D. C.; Biswas, S.; Zou, J.; Liu, Y.; Pandya, D.; Menachery, V. D.; Rahman, S.; Cao, Y.-A.; Deng, H.; Xiong, W.; Carlin, K. B.; Liu, J.; Su, H.; Haanes, E. J.; Keyt, B. A.; Zhang, N.; Carroll, S. F.; Shi, P.-Y.; An, Z. Nasal Delivery of an IgM Offers Broad Protection from SARS-CoV-2 Variants. Nature 2021, 595 (7869), 718–723. 10.1038/s41586-021-03673-2.

(2) Cuesta, Á. M.; Sainz-Pastor, N.; Bonet, J.; Oliva, B.; Álvarez-Vallina, L. Multivalent Antibodies: When Design Surpasses Evolution. Trends Biotechnol. 2010, 28 (7), 355–362. 10.1016/j.tibtech.2010.03.007.

(3) Braisted, A. C.; Wells, J. A. Minimizing a Binding Domain from Protein A. Proceedings of the National Academy of Sciences 1996, 93 (12), 5688–5692. 10.1073/pnas.93.12.5688.

(4) Ultsch, M.; Braisted, A.; Maun, H. R.; Eigenbrot, C. 3-2-1: Structural Insights from Stepwise Shrinkage of a Three-Helix Fc-Binding Domain to a Single Helix. *Protein Engineering*, Design and Selection 2017, 30 (9), 619–625. 10.1093/protein/gzx029.

(5) Khatri, B.; Pramanick, I.; Malladi, S. K.; Rajmani, R. S.; Kumar, S.; Ghosh, P.; Sengupta, N.; Rahisuddin, R.; Kumar, N.; Kumaran, S.; Ringe, R. P.; Varadarajan, R.; Dutta, S.; Chatterjee, J. A Dimeric Proteomimetic Prevents SARS-CoV-2 Infection by Dimerizing the Spike Protein. Nat. Chem. Biol. 2022, 18 (10), 1046–1055. 10.1038/s41589-022-01060-0.

(6) Kishimoto, S.; Nakashimada, Y.; Yokota, R.; Hatanaka, T.; Adachi, M.; Ito, Y. Site-Specific Chemical Conjugation of Antibodies by Using Affinity Peptide for the Development of Therapeutic Antibody Format. Bioconjug. Chem. 2019, 30 (3), 698–702. 10.1021/acs.bioconjchem.8b00865.

(7) Porter, R. R. The Hydrolysis of Rabbit γ-Globulin and Antibodies with Crystalline Papain. Biochemical Journal 1959, 73 (1), 119–127. 10.1042/bj0730119.

(8) Karageorgos, I.; Gallagher, E. S.; Galvin, C.; Gallagher, D. T.; Hudgens, J. W. Biophysical Characterization and Structure of the Fab Fragment from the NIST Reference Antibody, RM 8671. Biologicals 2017, 50, 27–34. 10.1016/j.biologicals.2017.09.005.

(9) Aschaffenburg, R.; Lewis, M.; Phillips, D. C.; Press, E. M.; Smith, S. G.; Sutton, B. J.; Mountford, C. W. Crystallographic Studies of Immunoglobulins: Crystallization of the Fc Fragment of Rabbit IgG with and without Cleavage of the Inter-Chain Disulphide Bridge. J. Mol. Biol. 1979, 135 (4), 1033–1036. 10.1016/0022-2836(79)90528-X.

(10) Dower, S. K.; Dwek, R. A.; McLaughlin, A. C.; Mole, L. E.; Press, E. M.; Sunderland, C. A. The Binding of Lanthanides to Non-Immune Rabbit Immunoglobulin G and Its Fragments. Biochemical Journal 1975, 149 (1), 73–82. 10.1042/bj1490073.

(11) Cole, J. L.; Lary, J. W.; P. Moody, T.; Laue, T. M. Analytical Ultracentrifugation: Sedimentation Velocity and Sedimentation Equilibrium; 2008; pp 143–179. 10.1016/S0091-679X(07)84006-4.

(12) Lahiri, P.; Verma, H.; Ravikumar, A.; Chatterjee, J. Protein Stabilization by Tuning the Steric Restraint at the Reverse Turn. Chem. Sci. 2018, 9 (20), 4600–4609. 10.1039/C7SC05163H.

(13) Tam, J. P.; Wu, C. R.; Liu, W.; Zhang, J. W. Disulfide Bond Formation in Peptides by Dimethyl Sulfoxide. Scope and Applications. J. Am. Chem. Soc. 1991, 113 (17), 6657–6662. 10.1021/ja00017a044.

(14) Horne, W. S.; Johnson, L. M.; Ketas, T. J.; Klasse, P. J.; Lu, M.; Moore, J. P.; Gellman, S. H. Structural and Biological Mimicry of Protein Surface Recognition by α/β-Peptide Foldamers. Proceedings of the National Academy of Sciences 2009, 106 (35), 14751–14756. 10.1073/pnas.0902663106.

(15) Schuck, P. Size-Distribution Analysis of Macromolecules by Sedimentation Velocity Ultracentrifugation and Lamm Equation Modeling. Biophys. J. 2000, 78 (3), 1606–1619. 10.1016/S0006-3495(00)76713-0.

